# Targeting Vitamin-D receptor (VDR) by a small molecule antagonist MeTC7 inhibits PD-L1 but controls THMYCN neuroblastoma growth PD-L1 independently

**DOI:** 10.1101/2020.08.16.252940

**Authors:** Rakesh K. Singh, KyuKwang Kim, Rachael B. Rowswell-Turner, Jeanne N. Hansen, Negar Khazan, Aaron Jones, Umayal Sivagnanalingam, Yuki Teramoto, Takuro Goto, Ye Jian, Nicholas Battaglia, Thomas Conley, Virginia Hovanesian, Naohiro Yano, Ravina Pandita, Leggy A. Arnold, Russel Hopson, Debasmita Ojha, Ashoke Sharon, John Ashton, Hiroshi Miyamoto, Nina F. Schor, Michael T. Milano, David C. Linehan, Scott A. Gerber, Richard G. Moore

## Abstract

Vitamin-D receptor (VDR) mRNA is enriched in malignant lung, ovarian and pancreatic tissues and showed poor prognoses. Calcitriol and stable or CRISPR-directed VDR upregulation increased PD-L1mRNA and protein expression in cancer cells in-vitro. A ChIP assay showed the binding of VDR with VDRE^PD-L1^. Stattic, a STAT3 phosphorylation inhibitor blocked calcitriol or VDR overexpression induced PD-L1 upregulation. MeTC7, a VDR antagonist developed by us, reduced PD-L1 expression on macrophages, ovarian, lung, breast, and pancreatic cancer cells in-vitro. In radiotherapy inducible PD-L1 model of orthotopic MC38 murine colon cancer, MeTC7 decreased PD-L1 surface expression, suppressed inflammatory monocytes (IMs) population and increased intra-tumoral CD69+PD1+CD8^+^T-cells. Intriguingly, MeTC7 reduced TH-MYCN transgenic neuroblastoma tumor growth without affecting PD-L1 and tumor immune milieu. In summary, Vitamin-D/VDR drives PD-L1 expression on cancer cells via STAT-3. Inhibiting VDR exhibited anti-checkpoint effects in orthotopic colon tumors, whereas PDL1-independent and anti-VDR/MYCN effects controlled growth of transgenic neuroblastoma and xenografted tumors.

**Summary:** Vitamin-D/VDR induces PD-L1 expression on cancer cells via STAT-3; and targeting VDR by a novel small molecule antagonist MeTC7 exhibits both anti-PD-L1 and anti-VDR/MYCN effects in tumor models.

## Introduction

Calcitriol, a VDR agonist, had exhibited anti-proliferative effects in experimental models of colon, breast, prostate and pancreatic cancer and hematologic malignancies (Giammanco et al., 2015). Phase-II/III clinical trials evaluating calcitriol showed lack of therapeutic responses and caused hypercalcemia (Beer et al., 2004, Trump DL et al., 2006). We demonstrate that VDR is overexpressed in malignant tissues of pancreatic, ovarian, and lung cancer compared to normal controls and correlates with poor prognoses. Calcitriol treatment transcriptionally upregulated PD-L1 gene and protein expression in a panel of cancer cells representing diverse tissue origins. VDR overexpression (stable and CRISPR directed) increased PD-L1 surface expression on ovarian and endometrial cancer cells. Interference with Stattic, a STAT3 phosphorylation inhibitor suggests that vitamin-D/VDR induced PD-L1 expression in pancreatic cancer cells is channeled through oncogene STAT-3. Calcitriol’s postulated role in tumor protection in colon, breast and pancreatic cancer contradicts Vitamin-D/VDR induced PD-L1 expression which is shown to promote tumor immune evasion (Sharma et al.,2015). In the context of Vitamin-D/VDR’s association with PD-L1 and oncogene STAT-3, it may appear that a VDR antagonist would be a better fitting tool to block VDR/PD-L axis than calcitriol which enhances expression of PD-L1 on tumor cells. In addition to the lack of therapeutic activities in agonists, progress in targeting VDR is hampered by the unavailability of pharmacologically pure VDR antagonists. Currently known VDR antagonists carry residual agonistic effects diminishing their pharmacologic value (Ishizuka et al., 2001). In this report, we demonstrate that Vitamin-D/VDR induced PD-L1 expression on tumor cells is routed through STAT-3, as Stattic, a phosphorylation inhibitor of STAT-3, abrogated calcitriol induced PD-L1 expression in BXPC-3 and PANC-1 pancreatic cancer cells. To target VDR, we describe the identification of a nuclear receptor (NR) selective and pharmacologically pure VDR antagonist, MeTC7, which inhibited PD-L1 surface expression on a panel of human and murine pancreatic, ovarian, breast and lung cancer cell-lines *in vitro*. Additionally, MeTC7 suppressed PD-L1 upregulation induced by radiotherapy (RT) in an orthotopic MC38-colorectal cancer model in mice (Gerber et al., 2013). In this model, MeTC7+RT treatment suppressed immune suppressive CD4^+^T cells (data not shown) and reduced the population of inflammatory monocytes (IMs) while increasing CD69^+^/PD-1^+^ CD8^+^T cell infiltration in the tumors. Intriguingly, MeTC7 treatment did not affect the immune niche of TH-MYCN transgenic murine model of neuroblastoma which involves VDR/MYCN/PD-L1 pathway (Weiss et al., 1997). Instead, MeTC7 blocked VDR/MYCN axis and controlled THMYCN driven neuroblastoma tumor growth without affecting expression of PD-L1 and other tumor immune markers in the tumors. In this study, association of Vitamin-D/VDR with PD-L1 and oncogene STAT-3 is shown and the effect of MeTC7 in the context of VDR/PD-L1 and VDR/MYCN/PD-L1 signaling is discussed.

## Results

### VDR was overexpressed in malignant tissues and correlated with reduced survival in pancreatic, lung, breast cancer and neuroblastoma patients

Mining of the publicly available microarray data at R2-genomics analysis and visualization platform (https://hgserver1.amc.nl/cgi-bin/r2/main.cgi) showed that compared to normal tissues, malignant tissues of the pancreas (p=0.0017), ovary (p=6.5e^-05^) and lung (p=5.4e-^08^) showed increased VDR mRNA expression (Figure-1A). Similarly, stroma of malignant ovarian tissues expressed elevated VDR mRNA expression compared to normal ovarian tissues (Figure-1A, p= 1.2e^-05^). VDR mRNA was also enriched in the tumor cells from recurrent neuroblastoma (Supplementary Figure-1). Kaplan-Meier survival of patients with lung and pancreatic cancer, grouped by the extent of VDR expression (from microarray data available at R2-genomics analysis and visualization platform and Human Protein Atlas (Uhlen et al.,2015), show that VDR mRNA enrichment significantly correlates with increased mortality (lung: p=0.00043, and pancreatic cancer: p=0.004) (Figure-1B-C). VDR mRNA enrichment also correlates with increased mortality in esophageal cancer and neuroblastoma (Figure-1 D-E). Increased mortality correlating with VDR enrichment was also found to be evident in breast cancer (p=0.011), glioma (p=0.0048), cervical cancer (p=0.055), liver cancer (p=0.048) and bladder cancer (p=0.05) (Supplementary Information-2). Although not statistically significant (p=0.09), the inverse association between VDR mRNA enrichment and mortality of ovarian cancer and SHH-B medulloblastoma patients was evident too (Supplementary information-2). Similar to VDR mRNA, VDR protein overexpression in malignant serous and endometrioid ovarian tissues compared to benign ovarian tissues was observed (Supplementary Figure-3: upper). Similarly, malignant tissues of cerebellum and fourth ventricle showed elevated VDR protein expression than normal cerebellum (Supplementary Figure-3: lower). We also analyzed whether the disease stage had any impact on the inverse association of VDR mRNA with decreased mortalities in cancer patients. Our analyses showed that among breast cancer patients with disease stage (IIa or IIb), VDR mRNA enrichment was not associated with increased mortalities (Supplementary Figure-4A-B). Among pancreatic cancer patients diagnosed with stage IIa and stage-IIb disease, VDR mRNA enrichment was strongly associated with increased mortalities (Supplementary Figure-4C-D). However, among ovarian cancer patients diagnosed with disease stage-II, increased VDR mRNA expression was not associated with increased mortalities either (p=0.19; Supplementary Figure-4E).

**Figure 1:**
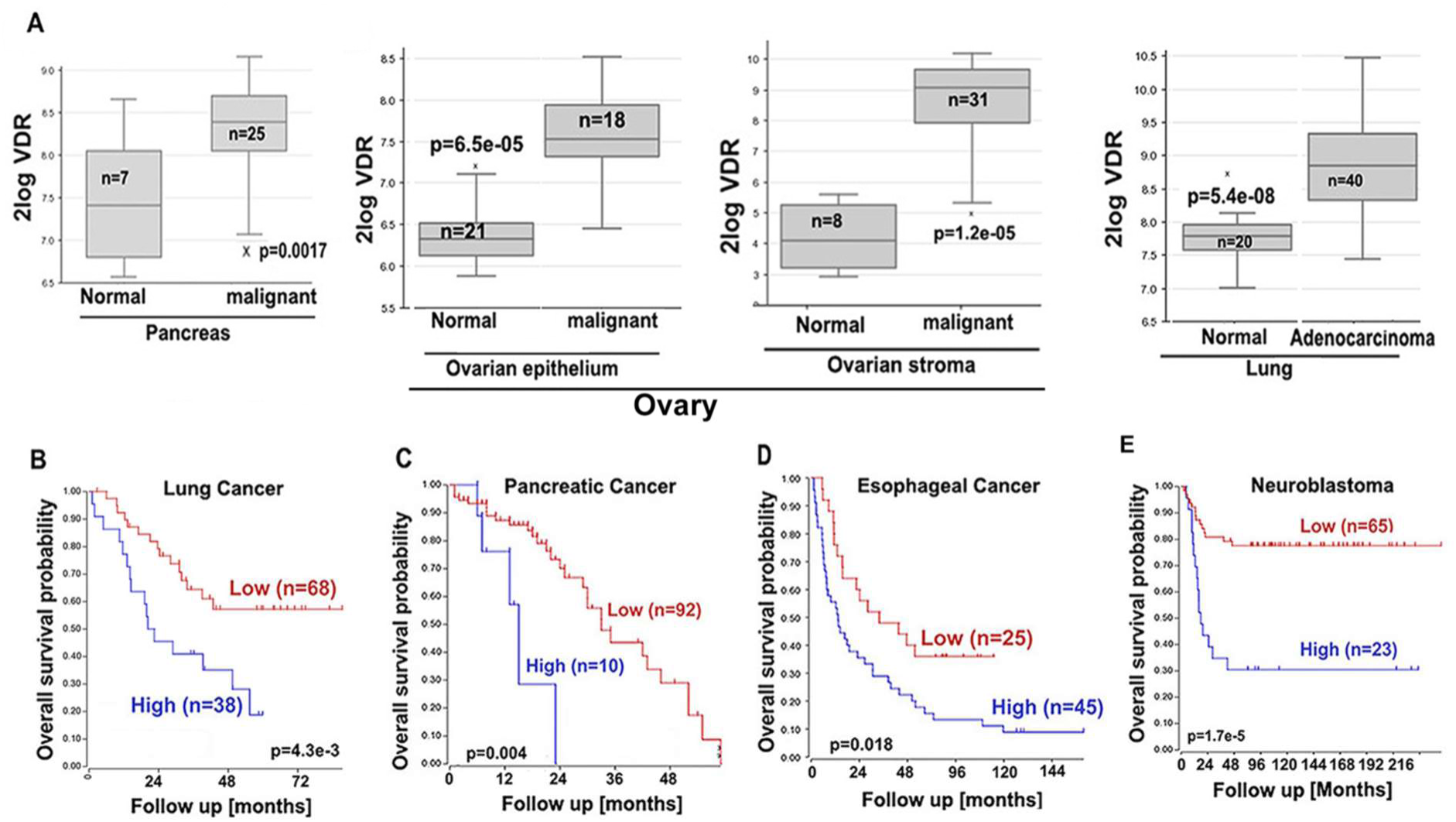
(**A**): Mining of microarray expression data available at R2 genomics analysis and visualization platform showed that VDR mRNA was enriched in the malignant tissues of pancreas, ovaries and lung compared to normal tissues. Stroma in ovarian malignant tissues also exhibited significantly increased VDR mRNA enrichment (third from left). VDR is also enriched in neuroblastoma cells from patients with recurrent/progressed disease (Supplementary Figure-1). (**B-E**): Kaplan Meier analyses of the pancreatic, lung, neuroblastoma, and esophageal cancer patients using the data and tools available at the R2 genomics and visualization platform or Human Protein Atlas show that VDR mRNA enrichment in lung, pancreatic and esophageal tumors and neuroblastoma correlate with decreased survival. Microarray data from breast, glioma, cervical, liver and bladder cancer also show statistically significant association of VDR mRNA enrichment with increased mortalities (Supplementary Figure-2). Although not statistically significant, microarray data from tumors of ovarian cancer (p=0.09) and SHH-β medulloblastoma (p=0.09) exhibited similar trend in the association of decreased survival of patients with increased VDR mRNA expression (Supplementary Figure-2).

### VDR overexpression increased surface expression of PD-L1 on cancer cells

Stably VDR overexpressing SKOV-3 ovarian cancer cell clones showed increased PD-L1 surface expression (Figure-2A). Immunoblot analyses of the total cell-lysates of VDR overexpressing SKOV-3 cell clones showed increased PD-L1 expression compared to pCMV (null vector) transfected control cells (Figure-2B, Left). Conversely, stable VDR knockdown in SKOV-3 cells resulted in reduced PD-L1 expression compared to null vector (Figure-2B, right). VDR’s impact on the upregulation of PD-L1 was further confirmed by CRISPR mediated VDR overexpression in ECC-1 endometrial cancer cell-line which also exhibited increased surface expression of PD-L1 (Figure-2C). RT PCR experiments confirmed that transcripts of both VDR and PD-L1 were upregulated in ECC-1 cells upon transfection with CRISPR plasmid (Supplementary Figure-5A-left: histogram; right: expression of VDR and PD-L1 transcripts).

**Figure 2:**
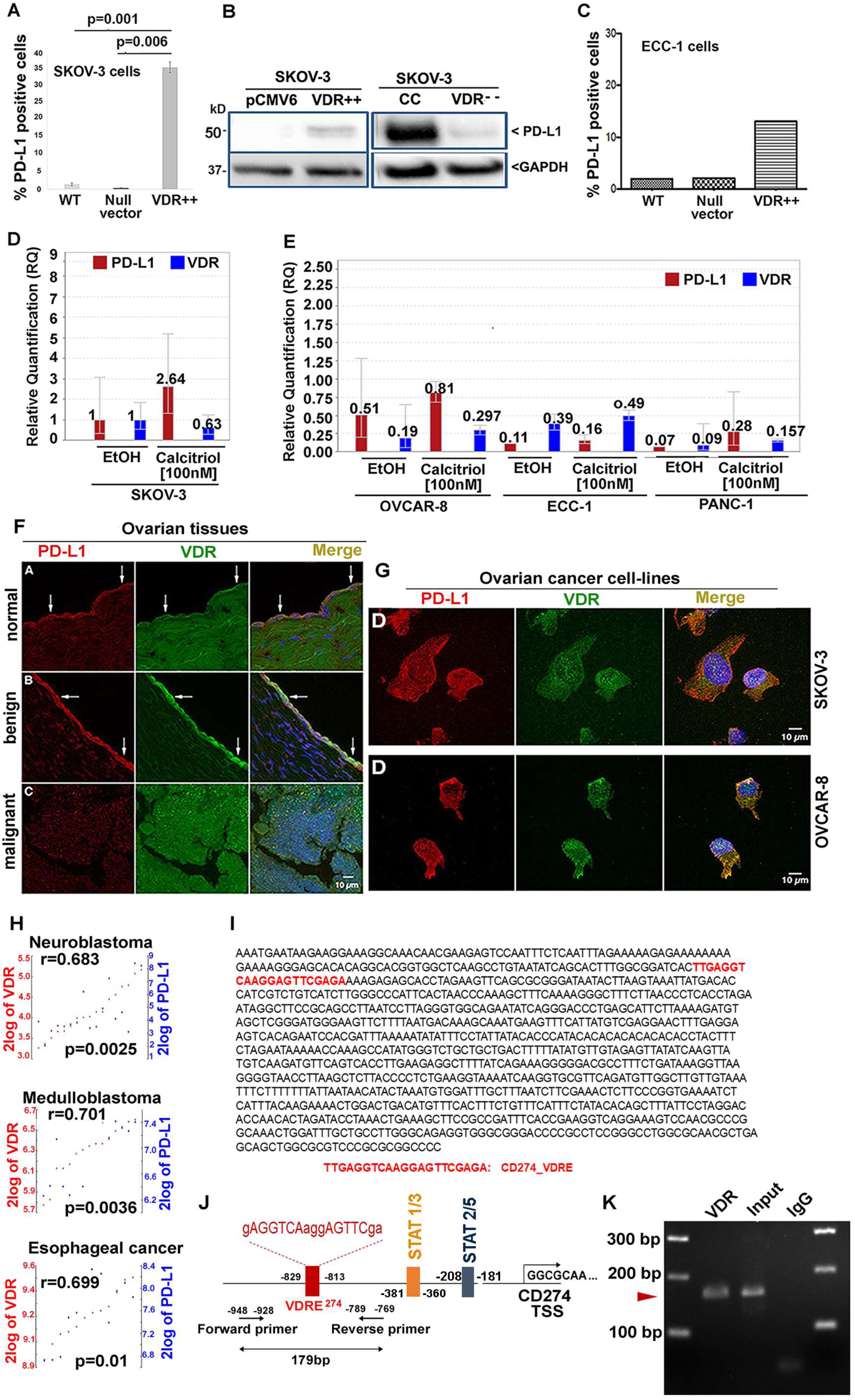
(**A**): Stable VDR overexpression in SKOV-3 EOC cells increased surface expression of PD-L1 compared to control and pCMV6 null vector. Surface expression of PD-L1 was estimated by flow-cytometry using a PD-L1 antibody (PDL1-BV605, Biolegend Inc, catalog number: 124321). Anti-IgG kappa beads (BD Biosciences. catalog number:552843) were used as negative controls. (**B**): Immuno-blot analysis of the total cell-lysates of VDR overexpressing SKOV-3 clone and null vector showed elevated expression of PD-L1 protein. Stable knockdown of VDR using shRNA showed decreased PD-L1 protein expression compared to control shRNA (CC) transfected SKOV-3 cells. Total cell-lysates were probed using PD-L1 antibody (Cell Signaling Technology, catalog number:13684S, dilution: 1:1000). PVDF membranes were stripped and re-probed for GAPDH as a loading control. (**C**): VDR upregulation via CRISPR activation plasmid increased PD-L1 surface expression on ECC-1 endometrial cancer cells. Both VDR and PD-L1 mRNAs were found to be co-upregulated (Supplementary Figure-5A) upon VDR overexpression. (**D-E**): SKOV-3 and OVCAR-8 (ovarian cancer), ECC-1 (endometrial cancer) and PANC-1 cells (pancreatic cancer) were treated with vehicle or calcitriol (100nM) for 48 hours. Relative VDR and PD-L1 mRNA expression in cells was analyzed by qpcr. Relative Quantification (RQ) values are shown on each bar. (**F-G**): VDR and PD-L1 exhibit co-localization in normal, benign and malignant ovarian tissues. Randomly selected tissue slides representing normal ovaries, benign ovaries and malignant serous ovarian tissues obtained from the pathology archives of Women and Infants Hospital of Rhode Island or (SKOV-3 and OVCAR-8 cell seeded slides) were processed, fixed and stained overnight with VDR primary antibody (Santa Cruz Biotechnology, catalog number:SC-9164, dilution: 1:500), washed and stained with source matched secondary (DyLight-488, Vector Laboratories, catalog number: DL-1488). Cells were washed and stained again with PD-L1 (Cell Signaling Technology, Mouse mAb catalog number: 29122, dilution: 1:1000) followed by staining with mouse secondary DyLight-594 antibody. DAPI containing mounting medium (Vectashield, Vector Laboratories, catalog number: H-1200) was applied and cover-slipped. Confocal images were recorded as described in material and methods section. Micron bars =10μm. (**H**): Analysis of microarray expression data available at R2-genomics analysis and visualization platform showed statistically strong correlation of VDR and PD-L1 in neuroblastoma, medulloblastoma and esophageal cancer patients. (**I**): Sequence of VDRE in PD-L1 promoter zone is shown. (**J**): Schema of VDRE, STAT/3 and STAT2/5 sequences and TSS of PD-L1 is shown. (**K**): ChIP assay using VDR (D6, sc-13133, Santa Cruz Biotechnology) antibody captured VDRE sequence in PD-L1 promoter. The sequence of VDRE primers (forward and reverse) are shown in Supplementary Information-9.

### Calcitriol upregulated PD-L1 in ovarian, endometrial and pancreatic cancer cells

Exposure of SKOV-3, OVCAR-8 (ovarian), ECC-1 (endometrial) and PANC-1 (pancreatic) cancer cells with calcitriol (100nM, 48 hours) showed increased PD-L1 mRNA expression when analyzed by qPCR (Figure-2D and E). VDR levels, except in SKOV-3 cells, were also increased compared in OVCAR-8, ECC-1 and PANC-1 cancer cells (Figure-2D-E). Similar to our findings, Dimitrov et al. had previously described Vitamin-D/VDR driven PD-L1 overexpression in human epithelial cancer cells (Dimitrov et al., 2017).

### VDR and PD-L1 showed co-localization in ovarian and medulloblastoma tissues

Confocal microscopy showed that VDR and PD-L1 co-localized in normal, benign, and malignant ovarian tissues (Figure-2F). In addition to ovarian tissues, ovarian cancer cell-lines SKOV-3 and OVCAR-8 also showed VDR and PD-L1 co-localization (Figure-2G). Averaged integrated optical density (IOD) values of the ten randomly selected tumor fields showed increased colocalization events in malignant and benign ovarian tissues than normal ovarian tissues (IOD values not shown). Medulloblastoma tissues also showed VDR and PD-L1 co-localization. (Supplementary Figure-5B). An immunoprecipitation experiment using VDR primary antibody showed enrichment of PD-L1 in SKOV-3 and OVCAR-8 (Supplementary Figure-5C-Left) and in DAOY and D283 medulloblastoma cells (Supplementary Figure-5C, Right). A two-gene (VDR and PD-L1) correlation analysis of microarray data available at R2-genomics analysis and visualization platform showed that VDR and PD-L1 exhibited strong correlation in neuroblastoma (r=0.683, p=2.5e^-03^), medulloblastoma (r=0.701, p=3.6e-03) and esophageal cancer (r=0.699, p=0.01) (Figure-2H) than pancreatic (Supplementary Figure-5D, r=0.255, p=0.08) and lung cancer (Supplementary Figure-5D, r=0.478, p=2.5e^-10^).

### Vitamin D receptor binds VDRE^CD274 (PD-L1)^ and directly regulates PD-L1 expression in PANC-1 cells

To determine whether Vitamin-D/VDR signaling up-regulates transcription of PD-L1, we searched for VDRE sequence in PD-L1 gene. In silico analysis of published promoter sequences of PD-L1 identified VDRE with 100% match (data not shown) between the base-sequences 829-813 (Figure-2I-J). We employed ChIP assays and conducted PCR of the bound sequences to confirm VDR binding to the VDRE sequence in PD-L1 promoter zone of PANC-1 cancer cells. Presence of VDRE sequence in anti-VDR antibody immune-precipitated chromatin sequence and the absence of it in the corresponding IgG control (Figure-2K) indicates that VDR binds VDRE^CD274 (PD-L1)^ and directly regulates PD-L1 expression in PANC1 cells and is suggestive of potential enhancer activity of VDR/VDRE.

### Stattic treatment abrogated Vitamin-D/VDR induced PD-L1 expression in cancer cell-lines

In silico analysis of the published PD-L1 promoter sequences revealed the presence of STAT1/3 and STAT2/5 in promoter zone of PD-L1 (Figure-2J). STAT/3 sequences were detected between base sequences (−381 and −360) whereas the STAT2/5 were detected between the sequences (−208 and –181). To capture a glimpse of the mechanism underlying the Vitamin-D/VDR mediated upregulation of PD-L1, the total cell-lysates of stable VDR overexpressing SKOV-3 cells were immunoblotted and probed with STAT-3 (phosphorylated and total). Our western blot analyses showed that stable VDR upregulation in SKOV-3 cells led to upregulation of PD-L1 and phosphorylated STAT-3 without affecting the expression of GAPDH (internal control) (Figure-3A). A treatment with Stattic (STAT-3 phosphorylation inhibitor) reduced PD-L1 expression in a stable VDR overexpressing SKOV-3 cell clone (Figure-3B). Mining of microarray expression of TH-MYCN tumors showed strong correlation in VDR and STAT-3 expression (r=0.919, p=1.19e^-83^) (Figure-3C). Similar strong correlation was also observed in pancreatic cancer microarrays (r=0.553, p=9.18e^-18^) (Figure-3D). A dose-dependent PD-L1 protein expression was observed upon treatment with calcitriol in BXPC-3 pancreatic cancer cells (Figure-3E). A 6-hour pretreatment with Stattic (500nM) in serum free medium blocked calcitriol induced PD-L1 expression in BXPC-3 (Figure-3F). Similarly, in PANC-1 pancreatic cancer cells, pretreatment with Stattic blocked increased transcriptional (Figure-3G) and protein expression (Figure-3H) of PD-L1 cell suggesting that calcitriol induced PD-L1 upregulation is STAT-3 dependent.

**Figure 3:**
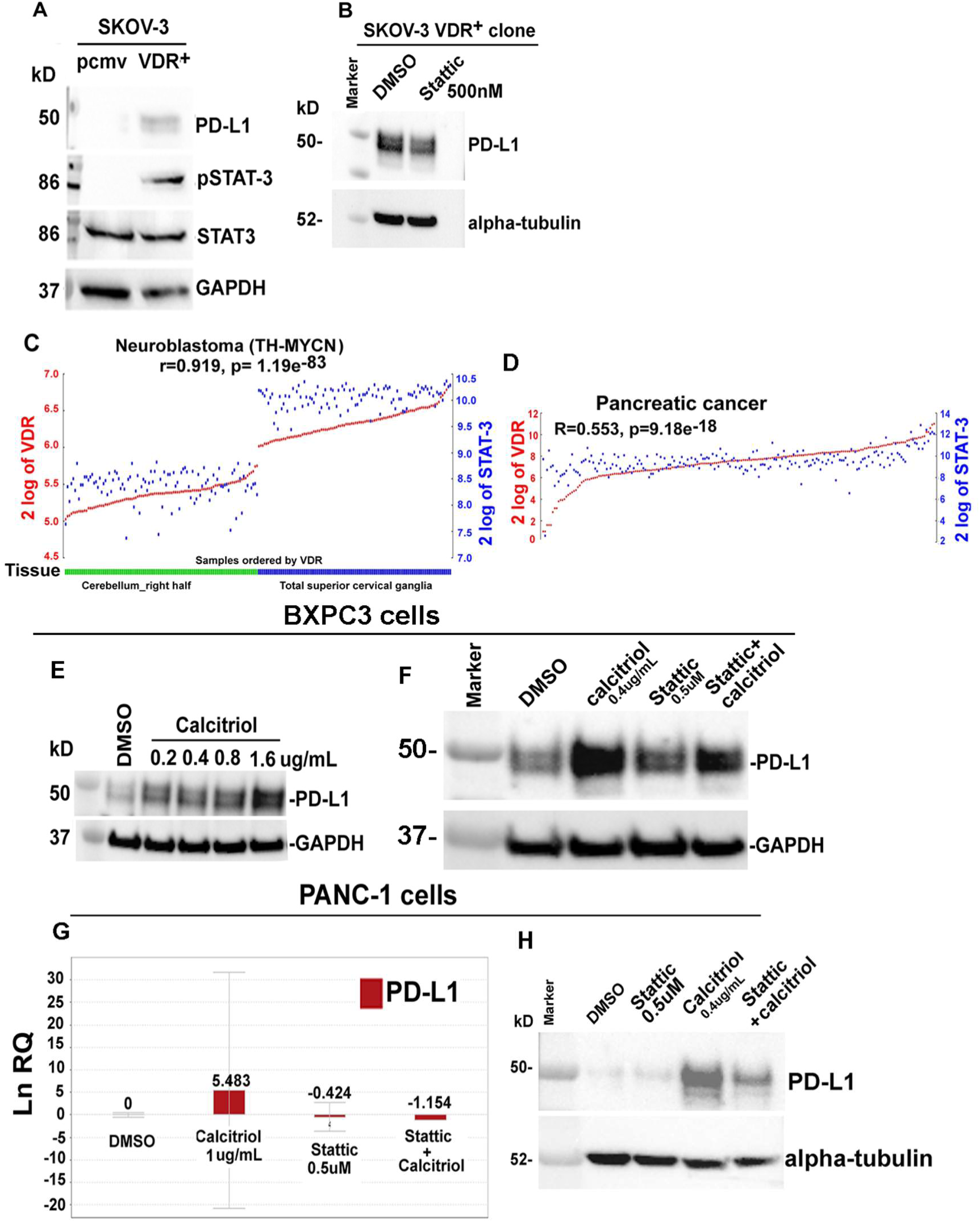
(**A**): Immunoblotting of VDR overexpressing SKOV-3 cell lysates showed increased PD-L1 and STAT-3 phosphorylation (Cell Signaling Technology, catalog number:9145). Total STAT3 (Cell Signaling Technology, catalog number: 30835) protein level was not altered. (**B**): Treatment with Stattic (500nM, 24 hours, STAT-3 phosphorylation inhibitor) reduced PD-L1 expression in a stable VDR overexpressing SKOV-3 cell-lines. (**C**): Mining of a neuroblastoma microarray database (TH-MYCN, R2-genomics analysis and visualization platform) exhibited strong correlation of VDR expression with STAT-3 expression (r=0.919, p=1.19e^-83^). (**D**): Mining of pancreatic cancer database also exhibited strong correlation of VDR and STAT-3 (r=0.553, p=9.8e^-18^). (**E**): Calcitriol (0-1.6ug/mL) dose-dependently increased PD-L1 protein expression in BXPC-3 pancreatic cancer cells. GAPDH levels were not altered. (**F**): Stattic (0.5nM) pre-treatment (6hrs) blocked calcitriol (0.4μg/mL) induced PD-L1 upregulation in BXPC-3 cells. GAPDH expression was not altered. (**G**): A 6hrs pretreatment with Stattic (0.5nM) blocked calcitriol (0.4ug/mL) induced PD-L1 mRNA expression in PANC-1 cancer cells. Naïve or treated cells were processed, and mRNA expression was analyzed by qPCR. (**H**): A 6hrs pretreatment with Stattic (0.5nM) blocked calcitriol(0.4μg/mL) induced PD-L1 protein upregulation in PANC-1 pancreatic cancer cells. Membranes were stripped and re-probed for α-tubulin expression, which as a loading control was not altered.

### Stable VDR knockdown reduced growth of SKOV-3 xenograft tumors

To determine the impact of VDR on tumor growth, we implanted VDR stably overexpressing clones (C12), VDR knocked down clones (C20) in nude mice subcutaneously along with cohorts implanted with wild-type SKOV-3 cells, cc clones (control to VDR under expressor C20 clones) and pCMV cell clones (control to VDR overexpressor cells). The growth of tumors was monitored by measuring the longest tumor diameter. As shown in the Supplementary Figure-6, compared to wild-type SKOV-3 cells and stable shRNA scrambled control, stable VDR knockdown SKOV-3 cells formed significantly slower tumor growth (wild-type vs C20: p= 0.0004; cc clone vs C20: p=0.0062). Surprisingly, C12 (VDR overexpressor clone) did not form an aggressive tumor phenotype and the difference in the tumor size between wild-type SKOV-3 vs VDR overexpressor clone (C12) derived xenograft tumors was insignificant (Supplementary Figure-6).

### MeTC7 is a NR selective VDR antagonist

MeTC7 (Figure-4A) (method of synthesis and characterization are shown in Supplementary Information-7) showed potent VDR inhibition (IC_50_ = 1.86 μM) (Figure-4B, left) in a fluorescence polarization (FP) assay performed using VDR-LBD and SRC2-3 Alexafluor 647. Fluorescence polarization studies showed that MeTC7 was void of any VDR agonistic activity (Figure-3B, right). In a cell-based transactivation assay, MeTC7 inhibited VDR transactivation in the concentration range of 13.58 to 46.28 μM (Figure-4C). MeTC7 showed similar activity as the calcitriol derivative (Cal-DT). In contrast, 7-dehydrocholesterol (7DHC), the precursor to Vitamin-D biosynthesis showed inhibition of VDR transactivation only at concentrations above 100uM (Figure-4C). It has been shown that the lack of NR selectivity is a major challenge with existing NR modulators (Burris et al., 2013). MeTC7 treatment did not show either agonistic or antagonistic effects on PPARγ (Figure-4D, left and right), a structurally homologous NR in the VDR family, concentration at or below 100 μM.

**Figure 4:**
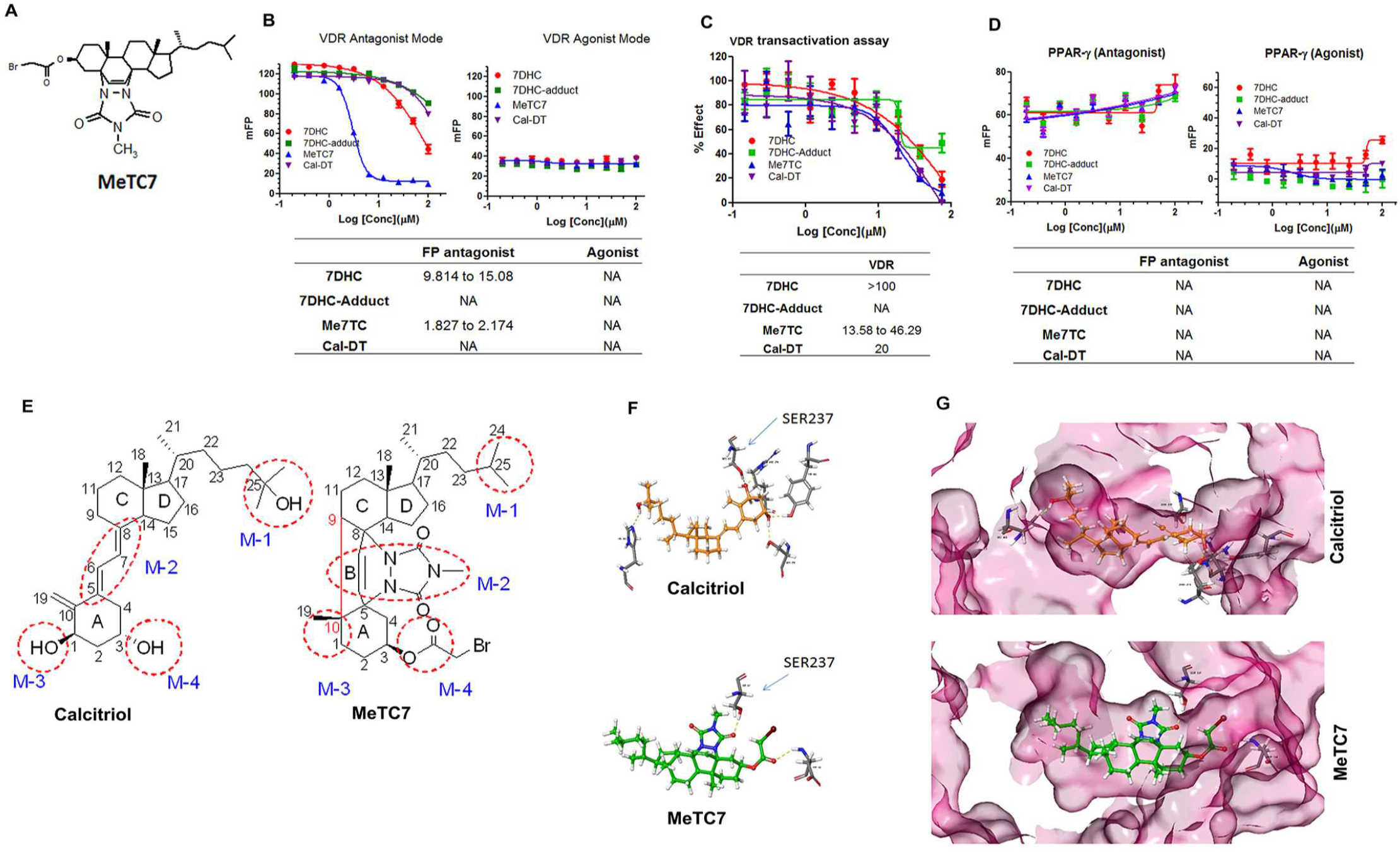
(**A**): Chemical structure of MeTC7. (**B**): Fluorescence polarization (FP) assay showed VDR antagonistic effect (left) without any residual agonistic effects (right). (**C**): MeTC7 treatment inhibited VDR transactivation in a cell-based assay. Interestingly, a calcitriol-adduct (Cal-DT) also showed similar inhibitory effect on VDR. **D**) MeTC7 did not exhibit any agonistic or antagonistic effects against PPAR-Y, a nuclear receptor that shares high sequence homology with VDR. (**E**): For *in-silico* binding to VDR, calcitriol (upper, left) and MeTC7 (lower, left) were marked into four structurally relevant zones (M1-M4). (**F**): Key interactions of both calcitriol (upper) and MeTC7 (lower) with Ser237 are shown. Snapshots of calcitriol (upper) and MeTC7 (lower) inserted into VDR-ligand binding domain (LBD) are shown. MeTC7 due to its structural bulk distorts VDR-LBD largely.

### In Silico studies identified the critical interactions of MeTC7 with VDR-ligand binding domain

*In silico* studies were performed to determine the interactions of MeTC7 with VDR-ligand binding domain (VDR-LBD) residues to elucidate the structural mechanism behind antagonistic characteristics of MeTC7. The crystal structure of VDR-LBD (IDB1) co-crystalized with 2-α-(3-hydroxy-1-propyl) calcitriol (Rochel et al., 2000) was used for docking studies (Figure-4 E-G). Our docking studies indicate that A-ring subsite (TYR143 and ARG 274 interaction with 1,3-OH of A-ring of calcitriol) was lost when MeTC7 bound with VDR (Figure-4F). MeTC7 interacts with a strong H-bonding with a backbone of ASP144 and SER 237 in the ring-A subsite (Figure-4F). MeTC7 utilizes only hydrophobic residues in the 25-OH-subsite, whereas calcitriol uses a hydrophilic interaction (SER306 & HIS305) as well. Calcitriol may be visualized to consist of a ring A, a conjugated linker, ring C and D along with a flexible chain as shown in Figure-4E (left). The simultaneous interactions mediated by C1, C3 and C25-OHs are crucial for super agonistic behavior of calcitriol. Synchronized interaction is possible only if correct spacing exists between the hydroxyl groups that are achieved by proper folding within the molecular structure of calcitriol to favor the correct orientation of hydroxyl group. Structurally, MeTC7 (Figure 4E-right) is larger in size and a conformationally rigid system compared to calcitriol. To handle this large and rigid molecule, the induced fit docking strategy (Xu et al., 2013) was implemented that revealed the possible binding mode of MeTC7 with VDR-LBD. The binding modes of MeTC7 are shown in Figure-4F in comparison with calcitriol. The major binding motifs (M1 to M4) in MeTC7 have been altered causing antagonistic effects. The shortening and removal of the C25-OH group near M1 motif of MeTC7 loses interaction with HIS 305. HIS 305 along with ARG 274 residues play a crucial role in determining the agonistic behavior of calcitriol and mutations at these residues leads to antagonistic effects (Mizwicki et al.,2009). The removal of conjugated linker system followed by shortening distance (2.6 Å) in MeTC7 in comparison to calcitriol (3.7Å) between ring A and C plays a leading factor for antagonistic behavior of MeTC7. Thus, the strategically placed triazolidine-dione moiety and the hydrophobic N-methyl group occupy the similar spatial position within VDR as was engaged by hydrophilic C1-OH of calcitriol. As a result, MeTC7 loses H-bonding with Arg 274, instead uses the carbonyl group to interact with SER 237 to retain stronger binding. Further, the conjugated diene linker of calcitriol enters tightly into the hydrophobic cavity of the VDR and agonize the system (Figure-4G-upper). In contrast, triazolidine-dione moiety of MeTC7 increases the volume of this VDR-LBD cavity and locks the conformational freedom of VDR due to deeper binding (Figure-4G-lower).

### MeTC7 reduced surface expression of PD-L1 *in vitro*

Immunoblotting of total cell lysates of OVCAR-8 (ovarian cancer) and NCI-H460 (lung cancer) cells treated with MeTC7 showed decreased PD-L1 expression (Figure-5A). Next, flow cytometry was employed to estimate the drug effect on the surface expression of PD-L1 on a panel of cancer cells after 48 hours of MeTC7 treatment. The panel of the cells included ovarian (ES2), pancreatic (PANC-1 and KCKO), lung (H1975) and murine breast cancer (E0771) cells (Figure-5B-G). Flow cytometric analysis of viable cells selected by live-dead staining demonstrated that MeTC7 treatment resulted in reduced PD-L1 surface expression at concentrations between 100-500 nM compared to IgG microbeads (Figure-5B-F). Both human (PANC-1, ES2, H1975) and murine cancer cells (KCKO and E0771) exhibited a decrease in PD-L1 surface expression at 100 nM except E0771 breast cancer cells, which required 500 nM treatment to exhibit PD-L1 reduction on the cell surface (Figure-5F). Similarly, MeTC7 (100 nM) treated macrophages showed lower PD-L1 surface levels compared to vehicle when cells were interrogated by flowcytometry (Figure-5G).

**Figure 5:**
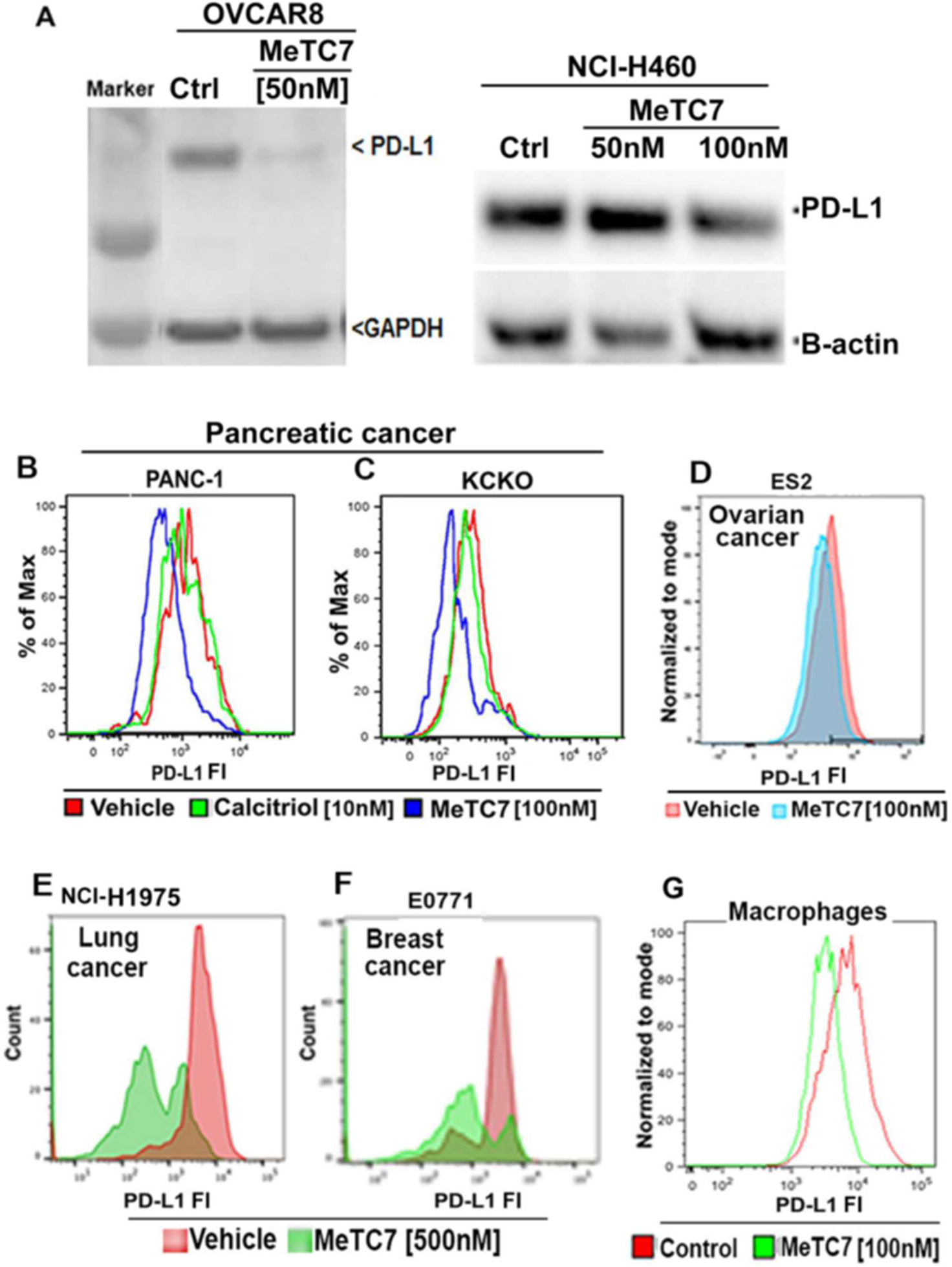
(**A**): MeTC7 (50-100nM, 48 hours) treatment reduced total PD-L1 levels in OVCAR-8 and NCI-H460 cells. The total cell lysates of vehicle or MeTC7 treated OVCAR8 and NCI-H460 cells were analyzed by immunoblotting techniques using PD-L1 monoclonal antibody (Cell Signaling Technology, catalog number:13684S, dilution: 1:1000). Either GAPDH or β-actin antibodies were used to assess the loading uniformity. (**B-D**): MeTC7 (100nM) treatment for 48 hours reduced surface expression of PD-L1 on human pancreatic cancer cell-line (PANC-1, **B**) murine pancreatic cancer cell-line (KCKO, **C**) and clear cell ovarian carcinoma cell-line ES2 in vitro (**D**). Calcitriol was used as a control. (**E-F**). Non-toxic dose of MeTC7 (500nM) reduced surface expression of PD-L1 expression on H1975 (human lung cancer cell-line) (**E**) and murine breast cancer cell-line (**F**). (**G**): Similarly, MeTC7 reduced PD-L1 surface expression on macrophages derived from CLL patient donated monocytes. The donor monocytes were differentiated by treatment with MCSF for 48 hours prior to treatment with MeTC7.

### MeTC7 inhibited radiotherapy induced PD-L1 expression in vivo

Next, we tested if MeTC7 treatment could reduce PD-L1 expression in an in vivo inducible PD-L1 activation model. It has been shown that radiotherapy (RT) induces PD-L1 expression on tumor cells in a MC38 cell derived syngeneic colorectal cancer model (Wu et al., 2016). To test if MeTC7 could prevent the RT-induced expression of PD-L1 on tumor cells, mice were injected with MC38 cells intramuscularly (i.m.) and tumor treated with 15 Gy local irradiation 7 days after inoculation. Vehicle control or MeTC7 was administered s.c. to mice daily on day 5 until endpoint (day 11) (Figure-6A, schema). Tumors were removed on day-11, dissociated into a single cell suspension, and analyzed by flow cytometry. Irradiated MC38 tumor cells exhibited a significant increase of PD-L1 expression when compared to unirradiated tumors (Figure-6B). Importantly, MeTC7 (25mg/kg) significantly reduced the surface expression of PD-L1 on tumor cells when compared to vehicle (Figure-6B). In addition, MeTC7, increased CD8+T-cell (CTLs) infiltration (Figure-6C). Flowcytometric analysis of the stained cells further showed that CD8+ T cell expressed increased CD69 and PD-1 activation markers when compared to both vehicle and RT+vehicle (Figure-6, D-E). Similarly, KLRG+T cells and inflammatory monocyte population in tumors were found to be significantly decreased in the MeTC7+RT group (Figure-6: F-G).

**Figure 6:**
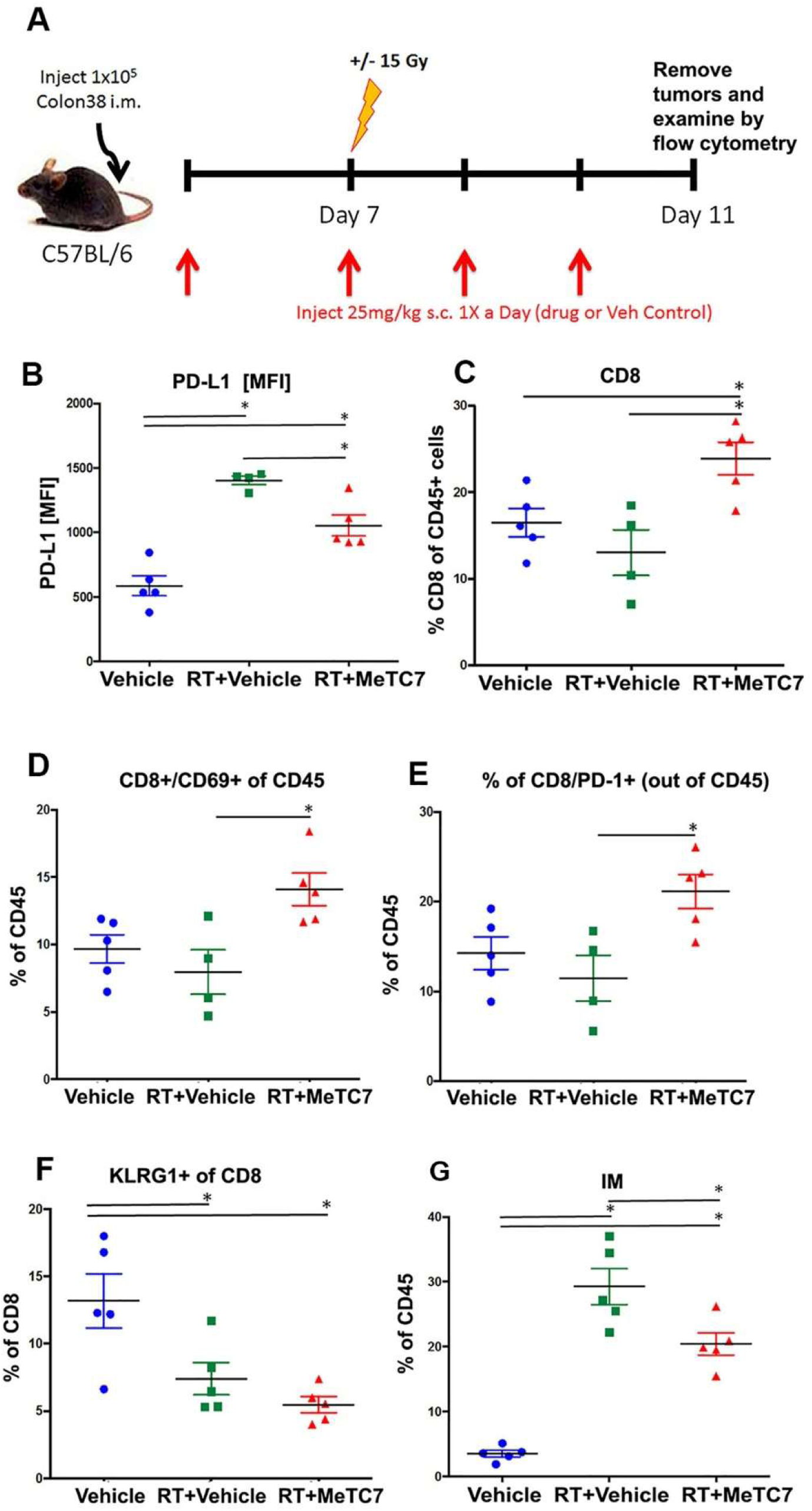
(**A**): Schema of evaluation of MeTC7’s activity against PD-L1 activated by RT in an orthotopic model of MC38 colorectal cancer in BL7 mice. Healthy MC38 cells were injected intra-muscularly in thighs of BL7 mice. On the day-7 mice (vehicle: n=5; MeTC7: n=5; RT: n=5; RT+MeTC7: n=5) were exposed to 15 Gy RT. MeTC7 at a dose of 25mg/kg subcutaneously was injected once daily until day 11 when mice were euthanized and tumors were isolated; fragmented to single cell suspension and analyzed by flow cytometry using a panel of murine antibodies. Analyses of the data showed that MeTC7 in combination with RT abrogated RT induced PD-L1 activation in tumors post RT (**B**), and (**C**) increased CD8+T-cell infiltration in tumors significantly compared to vehicle or RT+vehicle. (**D**): Combination of MeTC7 with RT increased activation marker CD69 on CD8+ cells and also PD-1 (**E**), another key CTL activation marker, compared to vehicle and vehicle+RT. (**F**): MeTC7+RT treatment reduced KLRG, a senescence marker of T cell. (**G**): MeTC7+RT treatment also reduced inflammatory monocytes (IMs). Catalog numbers of the antibodies used are listed in the Supplementary Information-9.

### MeTC7 blocked growth of TH-MYCN transgene driven spontaneous neuroblastoma

It has been shown that VDR drives MYCN, which, in turn induces PD-L1 expression on neuroblastoma tumor cells (Weiss et al., 1997). Therefore, we investigated whether MeTC7 can inhibit VDR, MYCN and its downstream PD-L1 and other tumor immune markers and control the tumor growth. TH-MYCN +/+ transgenic mice, which develop spontaneous neuroblastoma tumors that recapitulate human neuroblastoma disease (Kiyonari et al., 2015). H&E staining of tumors harvested from homozygous THMYCN+/+ mice (5.5 weeks old) showed strong presence of VDR, MYCN and TH antigens (Figure-7A). MeTC7 treatment reduced the viability of tumor spheres isolated from two independent homozygous THMYCN+/+ mice dose dependently (Figure-7B). Similarly, an MTS cell viability assay run on the tumor cells derived from three independent homozygous THMYCN+/+ mice showed that MeTC7 treatment was deleterious to tumor cell’s viability which decreased dose-dependently over three days of treatment monitoring (Figure-7C). Next the efficacy of MeTC7 against growing THMYCN+/+ homozygous tumors was tested in vivo. The response of the drug was monitored using Ultrasound imaging images and images were reconstructed to capture 3D tumor volume using inbuilt software. As shown in Figure-7D, MeTC7 (10mg/kg) reduced tumor growth when compared to vehicle. MeTC7 treatment Analysis of the tumor volume (p=0.033) (Figure-7B, middle) and end tumor weight analysis (p=0.053) (Figure-7D) demonstrated the tumor growth control mounted by MeTC7 treatment. To further validate the antitumor activity of MeTC7 against neuroblastoma, VDR and MYCN positive neuroblastoma xenograft tumors derived from SHSY5Y and Be2C neuroblastoma cells were treated with MeTC7 (10mg/kg, I.P., M-F). Tumor growth was monitored by measuring tumor volume (length x width^2^/0.5) periodically. Analyses of the tumor growth in the treatment group versus vehicle treated group showed that MeTC7 treatment reduced the growth of both SH-SY5Y (p=0.012, day:10^th^) and Be2C tumors (p=0.08, day-8th). Comparison of the tumor weights in the vehicle and drug treated Be2C tumors harvested at the conclusion of the experiment demonstrated the tumor growth control mounted by MeTC7 treatment (p=0.058).

**Figure 7:**
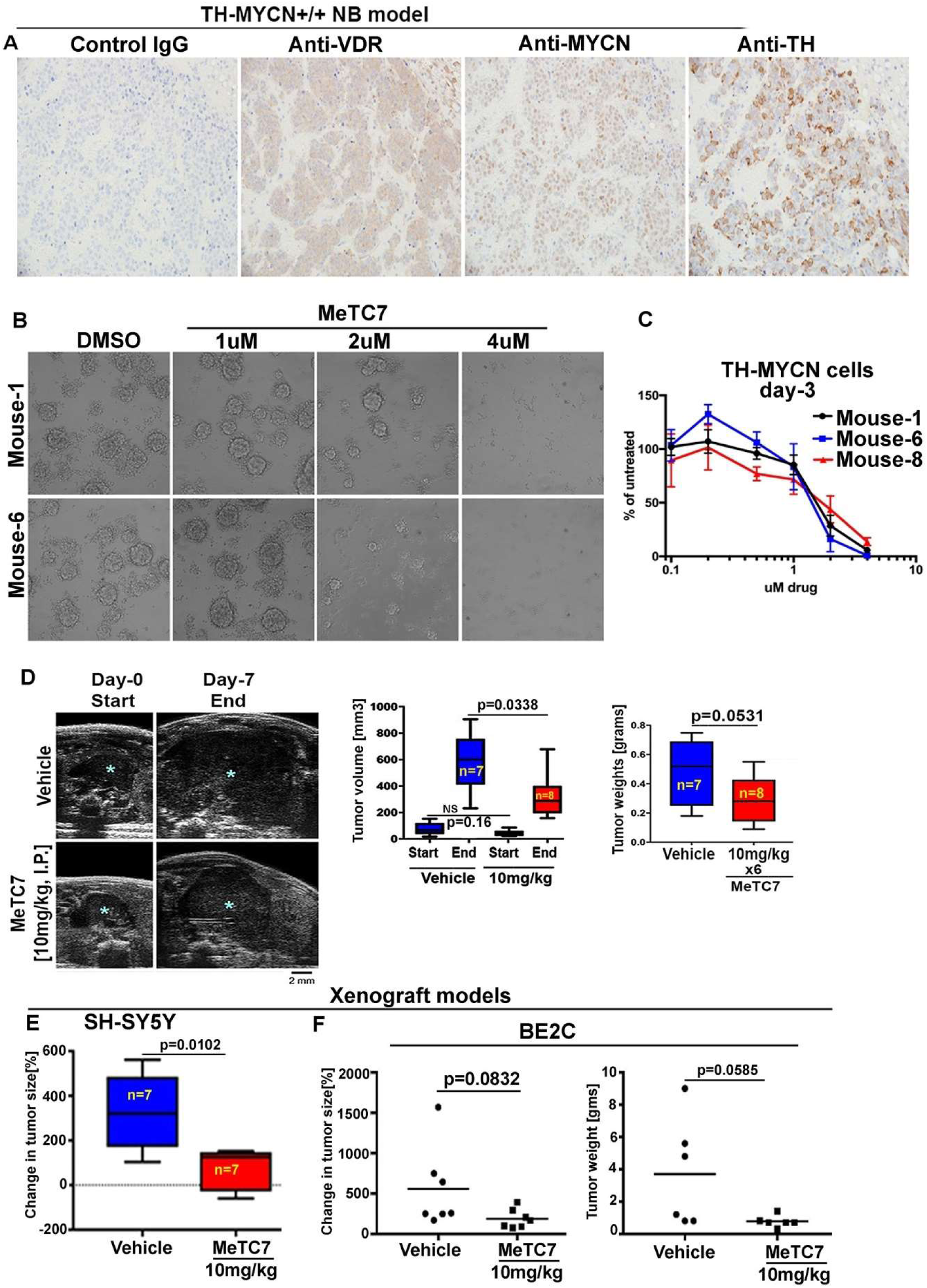
(**A**): Tumors from homozygous TH-MYCN (5.5 weeks old) mouse were isolated and H&E stained with IgG, VDR and MYCN antibodies. Tumors were also stained for the transgene-TH. Tumors showed strong expression of VDR, MYCN and TH. (**B**): MeTC7 treatment dose-dependently decreased the viability of murine neuroblastoma tumor spheres isolated from THMYCN mouse (tags: 1 and 6). Murine tumor sphere cells were grown in complete DMEM media and treated with varying doses of MeTC7. Representative images of the cells after 24 hours of treatment were recorded. (**C**): MeTC7 treatment reduced the viability of THMYCN tumor cells isolated from the three independent mice (tags: 1, 6 and 8) during three days of treatment. (**D**): Antitumor activity of MeTC7 in a TH-MYCN driven model of spontaneous neuroblastoma. (**D-left**): Compared to vehicle (upper, n=8), MeTC7 (lower, 10mg/kg, I.P., n=8) treatment reduced growth of THMYCN driven neuroblastoma in transgenic mice. Tumor burden was measured by an ultrasound imaging instrument. (**D-middle**): MeTC7 treatment dose-dependently reduced THMYCN tumor volume at the end of the treatment. (**D-middle**): Mice were euthanized and extracted tumors were weighed. (**E-F**): MeTC7 treatment reduced the growth of xenograft tumors derived from VDR high expressor neuroblastoma cells-lines (SH-SY5Y and Be2C). The response of MeTC7 against SHSY5Y xenograft tumors on day-10 of treatment is shown. Be2C cells (VDR++ and MYCN++) formed very aggressive tumors in mice and the response of the drug on day-8^th^ of the treatment is shown.

## Discussion

VDR enrichment and associated increase in mortalities among patients diagnosed with pancreatic, ovarian, lung and breast cancer and neuroblastoma; and immune checkpoint inhibitor ligand PD-L1 and oncogene STAT3 contradicts historically pursued anti-proliferative functions of Vitamin-D/VDR (Beer et al, 2004 and Trump et al, 2006). Association of Vitamin-D/VDR with PD-L1 and STAT-3 was established by stable or CRISPR directed VDR expression, co-localization of VDR with PD-L1 in ovarian cancer and medulloblastoma tissues and cells, immunoprecipitation of PD-L1 in ovarian cancer (SKOV-3, OVCAR-8) and medulloblastoma (DAOY, D283) cells via a VDR monoclonal antibody; and strong correlation (r>0.68) of VDR with PD-L1 and STAT-3 in neuroblastoma, pancreatic cancer and other cancer patient’s microarray data (Figure-1H, Supplementary Figure-5D), STAT-3 inhibitor mediated PD-L1 expression inhibition, and a ChIP assay which showed binding of VDR with VDRE^CD274^ and the in silico analyses that identified presence of VDRE and STAT-3 sequences in the promoter zone of PD-L1, which, taken together, suggest that aberrant VDR expression can play a role in tumor immune evasion and ensuing tumorigenesis. Likelihood that VDR is associated with tumorigenesis is further underlined by polymorphisms in melanoma, lung, pancreatic, prostate, ovarian and colorectal cancers (Gandini et al., 2014; Mocellin et al., 2008; Lei et al., 2013) that negatively impacts response to chemotherapy, overall survival, and time to disease progression (Schultheis et al., 2008, Woo et al., 2012); and VDR enrichment seen in premalignant and early malignant lesions (Agic et al., 2007) and the VDR mediated epigenetic corruption to block anti-proliferative genes in cancer cells (Abedin et al., 2006), suggesting that VDR inhibition should be the preferred approach to control malignancies than the historical approach of activating VDR via calcitriol or its analogs. VDR inhibition can control tumor growth was proven by reduced growth of VDR knocked down SKOV-3 tumors in vivo compared to scrambled vector and wild-type SKOV-3 controls (Supplementary Figure-6). However, the progress in targeting VDR has remained hampered by both, the lack of efficacy and hypercalcemia linked to VDR agonist’s use in phase-II/III clinical trials, and the unavailability of pharmacologically pure VDR antagonists. Literature described VDR antagonists carry residual agonistic effects that hinder their pharmacologic use in signaling and treatment studies (Ishizuka et al., 2001). Through our approach of chemical modification of vitamin-d related secosteroidal scaffold that started with the development of earlier classes of VDR antagonists (MT19c) (Moore RG et al., 2012), we have now developed MeTC7 which is more potent than MT19 in terms of VDR inhibition (Figure-4A). MeTC7, obtained by hetero-cyclization of 7-dehydrocholesterol (7DHC) with N-methyltriazoline-1,2,4-dione (MTAD) and subsequent esterification of the hydroxy-adduct with bromoacetic acid; is a pure VDR antagonist and is devoid of any agonistic activity and possesses high nuclear receptor selectivity (Figure-4B-D). We have noticed that heterocyclization of diene system in secosteroidal structure in Vitamin-D by Diel-Alder reaction with dienophiles like MTAD and PTAD removes the hypercalcemic liabilities of Vitamin-D and makes the resultant structures behave like antagonist without any residual agonistic liabilities. Thus, the development of MeTC7 addresses the need for a pharmacologically pure VDR antagonist that can be exploited to delineate the effects of VDR and its role in normal and malignant cells arising from its association with PD-L1 and oncogene STAT-3.

Not only did MeTC7 treatment decrease PD-L1 surface expression on pancreatic, ovarian, lung and breast cancer cells of human and murine origin *in vitro*; the radiotherapy inducible PD-L1 overexpression on MC38 colon cancer cell derived orthotopic tumor cells was significantly reduced *in vivo* also (Figure-6B). PD-L1 surface expression reduction on colon tumors was followed by increased CD8+T cell infiltration in tumors (Figure-6C). Further flow-cytometric interrogations showed that the tumor infiltrated CD8+T cells exhibited activation markers (CD69 and PD-1), suggesting that MeTC7 functions as a small molecule therapeutic checkpoint immunotherapy. Given the association of VDR with PD-L1 and oncogene STAT-3, through future studies we will interrogate whether MeTC7 mediated PD-L1 inhibition followed by downregulation of immune suppressive myeloid and inflammatory elements; and increased infiltration of activated CD8+T cells in radiotherapy treated colon tumors can lead to a meaningful control tumor growth; and assess whether MeTC7 could overcome the disadvantages associated with PD-L1/PD-1 targeting antibodies including poor solid tumor penetration and life-threatening immune adverse reactions (irAEs) (Lee et al.,2017, Gong et al., 2019). Similarly, while antibodies targeting PD-L1/PD-1 checkpoint axis have shown unprecedented improvement in tumor control and survival in certain malignancies (Coukas et al., 2011), limited eligibility of the patients (Zamarin et al., 2018) and life-threatening immune reactions (Topolian et al., 2012) remain major handicaps against their use. Antibodies also face *de novo* resistance; and generally, fail to penetrate solid tumor compartments, and are thus, unable to block secreted PD-L1 which was shown to promote resistance to immune checkpoint therapies in lung cancer (Gong et al., 2019). Advantages of small molecules include possibilities of the moderate adverse immune reaction events due to their shorter circulation life than antibodies, and the benefits of their abilities to penetrate solid tumor compartments to block the secreted form of PD-L1 which can prevent emergence of resistance against immune checkpoint therapies (Melaiu et al., 2017). While a small molecule inhibitor of PD-L1 has yet to be identified, a limited number of molecules that inhibit PD-L1 indirectly have been described (Prima et al., 2017; Lee et al., 2017). However, CA-170, a dual inhibitor of PD-1 and VISTA, is the only small molecule that is undergoing clinical trials for immuno-therapeutic treatment of malignancies (Lee et al., 2017).

Next, we attempted to determine whether targeting the VDR can generate a meaningful antitumor response in an aggressive neuroblastoma animal model that is driven by VDR. It has been shown that VDR regulates oncogenic transcription factor MYC which drives PD-L1 expression in neuroblastoma cells (Seuter et al.,2013; Salehi-Tabar et al., 2012; Veenstra et al, 1997). PD-L1 expression has been observed in tumor specimens of high-risk neuroblastoma patients (Chaudhary et al., 2015), and blockade of PD-1/PD-L1 in an animal model was shown to enhance the outcome of immunotherapy with anti-GD_2_ antibody (Ab) ch14.18/CHO (Rasmusan et al., 2012). Thus, VDR/MYCN/PD-L1 axis in neuroblastoma was found to be an appropriate model to test the effect of MeTC7 treatment on a TH-MYCN driven spontaneous neuroblastoma that closely recapitulates human neuroblastoma disease (Wang et al., 2015). Intriguingly, MeTC7 treatment which blocked MYCN and VDR expression in homozygous TH-MYCN tumor spheroids and exhibited reduced growth of TH-MYCN driven spontaneous neuroblastoma tumors in animals did not affect PD-L1 expression on tumors and other critical tumor immune niche markers (Supplementary Figure-8). Although, it has been suggested that pharmacologic inhibition of MYCN and MYC may be exploited to target PD-L1 to restore antitumor immunity in neuroblastoma (Melaiu et al, 2017), antagonizing the PD-1/PD-L1 axis failed to delay progression of established spontaneous tumors in the TH-MYCN mice (Mao et al, 2016). Promising antitumor effects in spontaneous THMYCN murine neuroblastoma and two xenograft models may suggest that targeting VDR/MYCN via MeTC7 can emerge as an effective approach to improve the neuroblastoma patient’s survival, utilizing, instead of VDR/PD-L1, the inhibitory effects on the VDR/MYCN pathway which may be a distinct and targetable mechanism of tumorigenesis in neuroblastoma.

## Materials and Methods

### VDR expression analyses

Survival analyses of patients diagnosed with pancreatic, lung, bladder, esophageal, bladder, neuroblastoma and other malignancies including the correlation of VDR with PD-L1 expression presented in this study (Figure-1, Figure-2H, and Supplementary Figure-1 and 2 and 5D) were generated by analyzing the microarray data available on the R2-Genomics Analysis and Visualization Platform (http://hgserver1.amc.nl) or on the Human Protein Atlas web site (https://www.proteinatlas.org/). Best system recommended cut-offs were selected.

### Methods of VDR activation and stable knockdown in ovarian cancer cells

For stable overexpression of VDR (Figure-2A), human VDR transcript variant 2 cDNA (OriGene, RC519628) was transfected into the SKOV3 cells using Lipofectamine 2000TM (Invitrogen) following the manufacturer’s instructions. Stably transfected cells were selected under the pressure of 500μg/mL of G418. Cells transfected with empty pCMV6-entry expression vector were used as control. For stable knockdown, SKOV-3 cells were transfected with human VDR shRNA (Santa Cruz Biotechnologies, SC-106692-SH) and the stably transfected cells were generated under 5 μg/mL of puromycin pressure. Cells transfected with a scrambled oligo vector (Plasmid-A, Santa Cruz Biotechnology, SC-108060) were used as control. Single cell cloning was done by limiting dilution method.

### CRISPR directed activation of VDR in ECC-1 endometrial cancer cells

ECC-1 cells were seeded at a density of 200,000 cells in 3 ml antibiotic-free standard growth medium per well (6-well plate) (Figure-2C and Supplementary Figure-5A). After reaching 70% confluency, cells were transfected with 2 μg of plasmid DNA. CRISPR overexpression plasmid DNA (Santa Cruz Biotechnology, sc-400171-ACT) was first added to OPTI-MEM medium (Gibco, catalog:31985-070) in a 1.5 mL microcentrifuge tube and allowed to sit for 5 minutes. The transfection reagent (Lipofectamine 3000; Invitrogen catalog: 1000022234) was added to OPTI-MEM medium in a microcentrifuge tube and allowed to sit for 5 minutes. The tubes containing plasmid DNA and transfection reagent were mixed together and allowed to sit for at least 20 minutes. Tubes were mixed well and 300 μL of plasmid DNA/transfection reagent was added to each well. Cells were incubated at 37°C, 5% CO_2_ for 24 hours, after which the media was replaced. Three days’ post-transfection cells were maintained under puromycin selection (1 μg/mL). Cells were kept in puromycin media until antibiotic-resistant clones grew. Once enough antibiotic-resistant cells were available, they were assayed for VDR protein and transcript levels via Western blotting and qPCR, respectively. Cells were assayed for PD-L1 surface expression via flow cytometry using PDL1-BV605 antibody (Biolegend Inc, catalog number:124321) using IgG beads (BD Biosciences, catalog number:552843) as negative control.

### Confocal and immuno-histochemical analyses of tumors

Immunohistochemical staining was performed on paraffin-embedded slides of ovarian tissue (normal, benign and malignant) specimens (thickness 5 μm) as described previously (Moore et al., 2014). Tissue sections were deparaffinized and rehydrated with serial ethanol dilutions of 100, 95 and 70%. Heat-induced antigen retrieval was then performed using DAKO Antigen Retrieval Solution for 20 min. Tissue sections were blocked with normal goat blocking serum (Vector Laboratories) for 60 min at room temp before incubating with primary antibodies for VDR (Santa Cruz Biotechnologies) and PD-L1 (Cell Signaling Technologies, 1∶200) in a humidified chamber overnight at 4°C. Secondary antibodies (DyLight 594 goat anti-rabbit IgG, Jackson Immuno-research Laboratories, INC. and Alexa Fluor 594 goat anti-mouse IgG at 1∶500, Invitrogen) were applied and incubated for 60 min for 1 hour at room temperature in the dark. Vectashield medium with DAPI (Vector Laboratories) was used to mount cover-slips for further analysis. Sixteen bit images were acquired with a Nikon E800 microscope (Nikon Inc., Mellville NY) using a 40× PlanApo objective. A Spot II digital camera (Diagnostic Instruments, Sterling Heights MI) was used to acquire the images (Figure-2E-F and Supplementary Figure: 3 and 5B). The cameras built-in green filter was used to increase image contrast. Camera settings were based on the brightest slide. All subsequent images were acquired with the same settings. Image processing and analysis was performed using iVision (BioVision Technologies, Exton, PA.) image analysis software. Positive staining was defined through intensity thresholding and integrated optical density (IOD) was calculated by examining the thresholded area multiplied by the mean. All measurements were performed in pixels. Confocal images were acquired with Nikon C1si confocal (Nikon Inc. Mellville NY.) using diode lasers 402, 488 and 561. Serial optical sections were performed with EZ-C1 computer software (Nikon Inc. NY). Z series sections were collected at 0.3 μm with a 40× PlanApo lens and a scan zoom of 2. The gain settings were based on the brightest slide and kept constant between specimens. Deconvolution and projections were done in Elements (Nikon Inc. Mellville, NY) computer software. Neuroblastoma tissues were fixed in 10% neutral buffered saline for several days, then dehydrated into paraffin using a Sakura VIP tissue processor and Sakura Tissue Tek 5 embedding center. Sections of 5-10μm in thickness were cut using a Leica RM2265 microtome for staining with either hematoxylin or eosin (H&E) or immuno-histochemical staining. Immuno-histochemical stains were performed using the GBI Polink-2 anti-rabbit HRP Plus Detection System (GBI International, D39) or Mouse-on-Mouse HRP-Polymer Bundle (BioCare Medical) and were counterstained with hematoxylin. Prior to primary antibody addition, sections were rehydrated, followed by 30 min antigen retrieval in sodium citrate buffer pH 6.0, and blocked of endogenous peroxidase with hydrogen peroxide. Primary antibodies used for immunohistochemistry were mouse rabbit anti-VDR (Abcam ab3508), mouse anti-VDR (Santa Cruz Biotechnology, sc-13133), mouse anti-MYCN (Santa Cruz Biotechnology, sc-53993), rabbit anti-tyrosine hydroxylase (TH) (Millipore AB152), normal rabbit IgG (Millipore 12-370), or normal mouse IgG (Millipore 12-371). Slides were visualized using an Olympus BX41 light microscope and imaged with an Olympus DP70 camera. Photographs were captured using CellSens digital software.

### Chromatin Immunoprecipitation (ChIP)

ChIP assay was performed using the Magna ChIP kit (Millipore, MA, USA) according to the manufacturer’s instructions with minor modification. PANC-1 cells were cross-linked with 1.0% formaldehyde for 10 minutes at room temperature. Sonications were done in nuclear buffer (four 30-s pulses, output 3.0, duty cycle 30% in ice with 120-s rest between pulses; Branson Sonifier 450). A fraction of the mixture of protein-DNA complex was used as “input DNA.” Soluble chromatin was immunoprecipitated with anti-VDRE antibody (D-6, Santa Cruz Biotechnology Inc) and normal mouse immunoglobulin G (IgG) (sc-2025, Santa Cruz Biotechnology Inc) directly conjugated with Magnetic Protein A beads. Immuno-precipitated DNA was eluted and reverse cross-linked, and then DNA was extracted and purified using a spin filter column. DNA samples were analyzed by PCR. PCR products were electrophoresed on 1% agarose gel, and ethidium bromide stained DNA was visualized by a Gel Doc XR+ (Biorad, Hercules, CA).

#### Synthesis of MeTC7

Synthesis of MeTC7 is described in Supplementary Information-7A. Structure of MeTC7 was confirmed by ^1^H, ^13^C and correlative NMR experiments (Supplementary Figure-7B).

#### Cell culture and cytotoxicity assay

TH-MYCN+/+ cell lines were derived by mechanical dissociation of tumors obtained from TH-MYCN homozygous mice (Chesler et al, 2011). These cells were maintained in RPMI 1640 media (Gibco, catalog number: 11875) supplemented with 20% heat-inactivated FBS, 10^−5^ mM 2-mercaptoethanol, 1 mM sodium pyruvate, and 1Xnon-essential amino acids (Gibco, catalog number: 11140076). SKOV-3, OVCAR-8, PANC-1, KCKO, DAOT, D283 and E0771 cells were grown in complete DMEM. ECC-1 and H1975 cells were grown in complete RPMI medium. MC38 murine colorectal cancer cells were grown in complete DMEM medium. ES2 was grown in McCoy’s 5A complete medium.

#### Determination of antagonistic properties of MeTC7 against VDR and PPAR-Y

Agonistic and antagonistic activity of MeTC7 against VDR and PPAR-Y was studied using a FP assay (Feau et al., 2009). This assay was conducted in black polystyrene plates (Corning Inc) using a buffer [25 mM PIPES (pH 6.75) 50 mM NaCl, 0.01% NP-40, 2% DMSO], VDR-LBD protein (0.1 μM), LG190187 (3 μM), and Alexa Fluor 647-labeled SRC2-3 (5 nM). Fluorescence polarization was detected after 1 hour at excitation and emission wavelengths of 650 nm and 665 nm, respectively. To determine the activity against PPARγ, PPARγ-LBD was expressed in BL21 (DE3) (Invitrogen), purified by affinity chromatography, and stored at −80°C in buffer (50 mM Tris (pH 8.0), 25 mM KCl, 2 mM DTT, 10% glycerol, 0.01% NP-40). For the assay, MeTC7 was serially diluted in DMSO and 100 nL of each concentration was transferred into 20 μL protein buffer (20 mM TRIS (pH 7.5), 100 mM NaCl, 0.01% NP-40, 2% DMSO, 10 nM DRIP2 (CNTKNHPMLMNLLKDNPAQD) labeled with Texas-Red maleimide, and 1 μM PPARγ-LBD) in the presence and absence of GW1929 (5 μM) in quadruplet using black 96 well plate (Costar, catalog number: 3658). The samples were allowed to equilibrate for two hours. Binding was then measured using fluorescence polarization (excitation 595 nm, emission 615 nm) using a Tecan M1000 plate reader. The experiments were evaluated using GraphPad Prism 5, and IC_50_ values were obtained by fitting the data to an equation (Sigmoidal dose-response-variable slope (four parameters). Values are given as the mean values of two independent experiments with a 95% confidence interval. The data were analyzed using nonlinear regression with a variable slope (GraphPad Prism).

#### Immunoprecipitation of VDR in ovarian cancer and medulloblastoma cells

Total cell lysates of ovarian (SKOV-3 and OVCAR-8) cells and medulloblastoma (DAOY and D283) cells were incubated with a primary VDR antibody (Santa Cruz Biotechnology, catalog number: VDR antibody (H-81): SC-9164) or a corresponding IgG antibody. The beads were washed repeatedly with washing buffer. The proteins were separated from the beads and analyzed by immunoblotting using primary antibodies against VDR and PD-L1. Similarly, the VDR antibody (Santa Cruz Biotechnology, catalog number: VDR Antibody (H-81): SC-9164) was utilized to immuno-precipitate PD-L1 from the total cell lysates of DAOY and D283 medulloblastoma cells. The immunoblots were probed with PD-L1 antibody (Cell Signaling Technology, rabbit mAb, catalog number: 13684).

#### Estimation of effect of MeTC7 treatment on surface expression of PD-L1 in cultured cancer cells

To determine the impact of MeTC7 treatment on total PD-L1 levels in cultured cancer cells, OVCAR-8 ovarian cancer cells and NCI-H460 lung cancer cells were treated with MeTC7 (50, 100nM) respectively for overnight. The total cell lysates were probed with the PD-L1 monoclonal antibody (Cell Signaling Technology, catalog number:13684S, dilution: 1:1000). GAPDH and β-actin were used as loading control (Figure-5A). To determine the impact of MeTC7 treatment on surface expression of PD-L1, cultured PANC-1 and KCKO (human and murine pancreatic cancer cells); ES2 (clear cell ovarian carcinoma cells). NCI-H460 (lung cancer), E0771 (murine breast cancer cells) and macrophage cells were treated with non-toxic concentrations (100nM or 500nM for NCI-H1975 and E0771 cells). Prior to treatment with MeTC7, the donor monocytes were differentiated by treatment with MCSF over a period of 48 hours before treatment with MeTC7 was carried out. The cells were processed and analyzed by flow cytometry to assess the PD-L1 levels on cells. Anti-mouse IgG antibody coated beads (BD Biosciences, catalog number: 552843) were used as the internal IgG control. Both PANC-1 and KCKO cells were treated separately with VDR agonist calcitriol as positive control.

#### Orthotopic colorectal cancer model of PD-L1 activation

MC38 tumor cells (1 x 10^5^) were injected intramuscularly in the left legs of female C57BL/6J mice. Sample size consisted of vehicle (n=5), MeTC7 (n=5), RT(n=5) and RT+MeTC7 (n=5). Mice were treated locally with RT 7 days after tumor cell injection using a 3200 Curie-sealed ^137^Cesium source that operates at roughly 1.90 Gy/min. Jigs were constructed and designed to specifically treat the tumor bearing leg with 15 Gy radiations (Gerber et al., 2013). This source and the collimators used are calibrated periodically to ensure equal distribution of radiation. Standard caliper was used to measure tumor growth. Tumor-bearing mice were administered 25 mg/ kg of MeTC7 or vehicle control (40% Hydroxypropyl-beta-cyclodextrin [Acros Organics] & solutol HS15 [Sigma] in sterile water) subcutaneously (s.c.) 1X/day starting 2 days before RT for the indicated amount of time. Experiment was carried out once.

#### Flow-cytometric analysis

Peripheral blood was collected from tail veins at various time points into tubes containing heparin (Hospira, Inc.). Tumors were removed 4 days post-RT and processed into single cell suspensions as previously described (Gerber et al., 2013). A total of 1 x 10^6^ tumor cells and 15uL of whole blood were blocked with Fc Block (clone 2.4G2) followed by staining with a cocktail of directly conjugated primary antibodies (Supplementary Information: 9) for 30 minutes. VDR overexpressing SKOV-3 and ECC-1 cells were processed and stained with PDL1-BV605 antibody (Biolegend Inc, Catalog number:124321). VDR expressing SKOV-3 and ECC-1 cells, tumor cells from mice and the peripheral blood cells were washed with 1 mL of PBS/1% BSA/0.1% azide, fixed with BD Cytofix/Cytoperm (BD Biosciences), and analyzed using a 12-color LSRII (BD Biosciences) and FlowJo software (Tree Star). For mice tumor and blood cells, the data is reported as percent of CD45+ events and normalized per milligram of tumor where indicated. Anti-mouse IgG antibody coated beads (BD Biosciences, catalog number: 552843) as internal IgG control.

#### Efficacy of MeTC7 in THMYCN transgenic neuroblastoma mice

TH-MYCN hemizygous mice (129X1/SvJ-Tg(TH-MYCN)41Waw/Nci) were initially obtained from the NCI Mouse Repository (strain code 01XD2) and maintained in a 129X1/SvJ background though cross-breeding with either wild-type 129X1/SvJ mice obtained from The Jackson Laboratory (stock number 000691) or other TH-MYCN hemizygous mice. TH-MYCN homozygous mice were identified through genotyping as previously described (Haraguchi et al., 2009). All mice were maintained on a breeder diet (Labdiet 5021) and tumor-bearing mice were further supplemented with Diet Gel® 67A (ClearH2O®). Mice in control (n=7), MeTC7 (10mg/kg, n=8) mice were treated intraperitoneally with indicated doses. Mice in control and MeTC7 (10mg/kg) group received six treatments in total, whereas the mice in MeTC7(100mg/kg) group were given just three treatments to see the effect of escalated drug dose on tumor burden (data not shown). Experiment was carried out once. The tumor burden in each mouse was estimated using ultrasound imaging instrumentation as described below. Portion of tumors from three homozygous THMYCN+/+ each in control and treatment group were broken into single cell suspension and tumor immune antigens were analyzed by flow cytometry as described above. The flow cytometry data is shown in Supplementary Figure-8.

#### Ultrasound imaging of the TH-MYCN mice

Tumors in vehicle/drug treated group were visualized by abdominal ultrasound using a Vevo 3100 Imaging System and MX550D transducer (FUJIFILM VisualSonics, Inc). Animals were anesthetized (1-3% isoflurane and oxygen mixture) and restrained on a heated stage with monitors for respiration and heartbeat. Ventral hair was removed with depilatory cream prior to monitoring with ultrasound probe. 3D volume measurements were carried out using Amira 6.1 software with a XImagePAC extension (FEI).

#### Xenograft animal model

SKOV-3 (wild-type), pCMV, CC and VDR stably overexpressing clones (C12) and VDR stably knocked clone (C20) (Supplementary Figure-6) and SHSY5Y and Be2C (Figure-6E-F) cells isolated from 70-80% confluent petri-dishes were spun down (1000 RPC, 5 minutes). Media was removed and cells (1million) were suspended in matrigel:serum free RPMI media mix (1:1) and implanted subcutaneously in the right flank of the nude mice. For SHSY5Y and Be2C cells, NSG mice were used. Prior to inoculation, NSG mice were shaved at the inoculation site using a clean shaving machine and skin was cleaned using alcohol swaps. After tumors became palpable, mice were treated with vehicle (n=5 for SKOV-3 clones and n=7 for SHSY5Y and Be2C each) and MeTC7 (n=7) intraperitoneally till tumor volumes [length x width^2^)/0.5] reached 2000mm^3^. Tumor sizes and animal weights were recorded every third day. Be2C cells formed highly aggressive tumors and treatment had to be stopped early due to tumor volume reached 2000mm^3^. Mice in the control and drug groups were euthanized and tumors were harvested. The % change in tumor growth and the difference in the tumor volumes and tumor weights in the treatment groups were compared to vehicle treated groups using Graph Prism software.

#### Statistical analyses

Statistical analyses of the tumor sizes between the control and MeTC7 treated group were determined using Graphpad Prism 7 software (GraphPad Inc.). Two tailed unpaired t-test was used to determine the difference between two groups. F-test was conducted to determine the size of the difference. Ordinary one-way ANOVA test was employed to determine the statistical difference between groups at multiple time points as the tumor treatment progressed. Brown-Forsynthe and Bartlett’s tests were performed to assess the statistical differences.

## Acknowledgement

JNH thanks Dr. Ron Wood and Dr. Deanne Mickelsen for ultrasound training/assistance. Nicole Romano PhD (Women and Infant’s Hospital of Rhode Island) is acknowledged for staining tumor slides for co-localization studies.

## Author contribution

RKS conceived the project, designed and synthesized the MeTC7 and organized the studies. SAG and YJ designed and completed the evaluation of MeTC7 in radiation induced PD-L1 activated colon cancer model. JNH designed and performed experiments related to TH-MYCN model under supervision of NFS. PD-L1 levels on stably overexpressing SKOV-3 cell or clones thereof were analyzed by RRT. YN developed VDR stably overexpressing and knockdown clones of SKOV-3. ALA designed and directed the fluorescence polarization assay to measure agonistic or antagonistic activity against VDR. MS in supervision of JA developed CRISPR activated ECC-1 clones and measured the PD-L1 levels. MS also measured PD-L1 levels on macrophages. RH performed extensive NMR experiments to establish the structure of MeTC7. TC conducted the flow cytometry studies on cells treated with MeTC7. Western blots and pcr experiments were done by AJ, NK, RP and KKK. RKS wrote the manuscript. TY and TG conducted ChIP assay. DL, MTM, JA, HM, NFS and RGM analyzed data, reviewed and edited the manuscript or contributed resources.

## Supplementary Figures

**Supplementary Figure 1:**
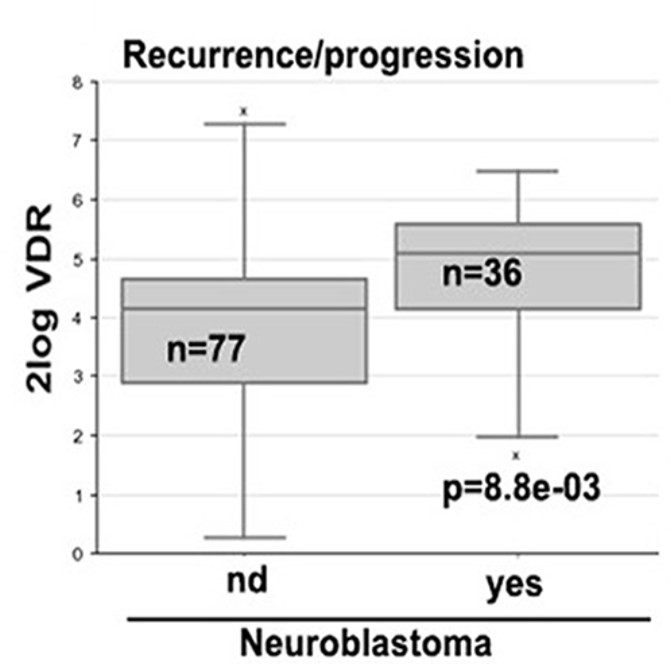
VDR mRNA is enriched in recurrent/progressed neuroblastoma. Microarray expression data available at R2 Genomics Analysis and Visualization Platform was analyzed using inbuilt tools and the system generated best expression cut-offs were selected.

**Supplementary Figure 2:**
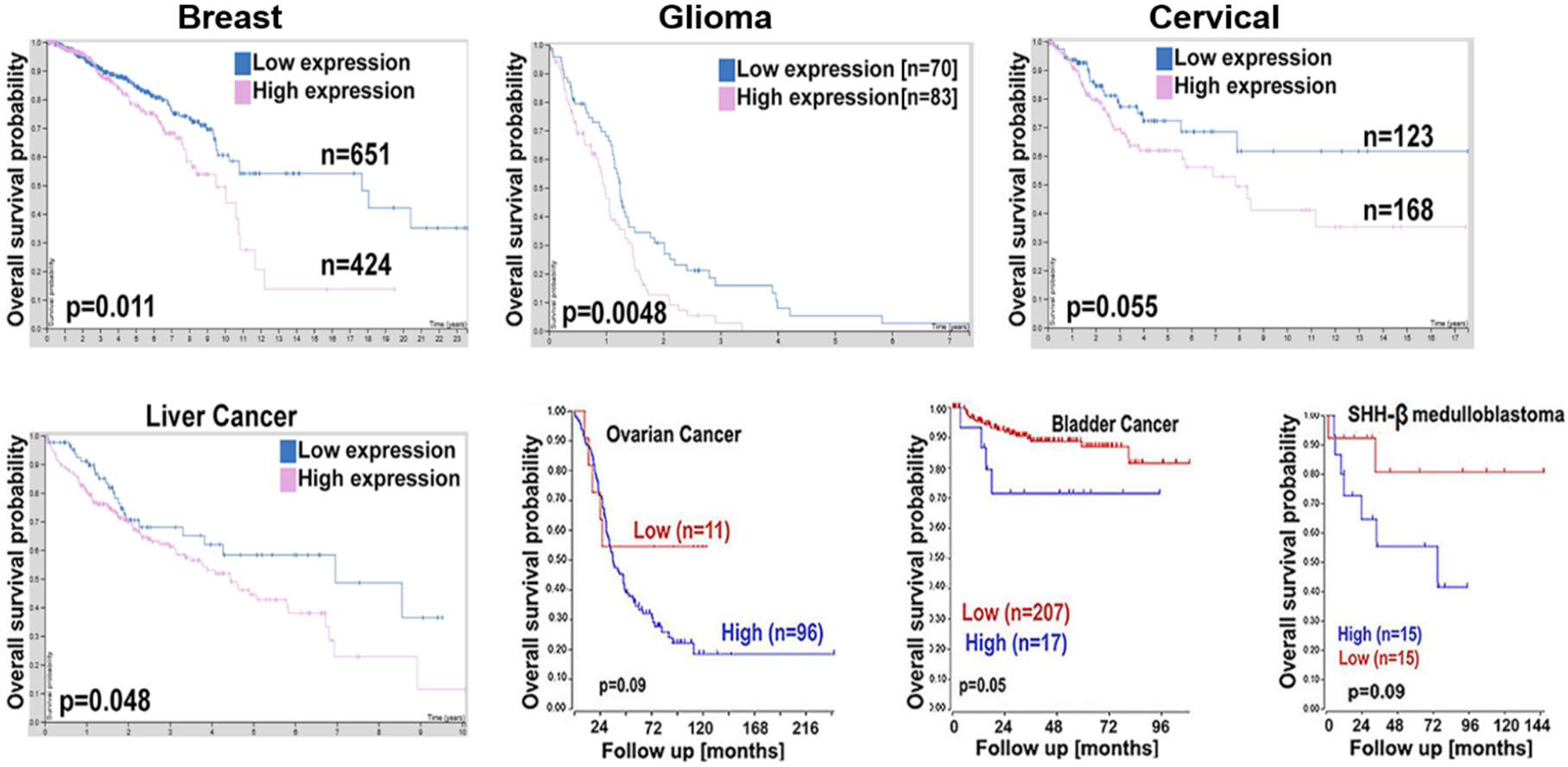
Kaplan Meier survival analyses of patients with bladder cancer, ovarian cancer and SHH-B medulloblastoma, show that VDR mRNA overexpression correlates with reduced survival. Microarray expression data available at R2 Genomics Analysis and Visualization Platform or Human Protein Atlas were analyzed using inbuilt tools and the system generated best expression cut-offs were selected.

**Supplementary Figure 3:**
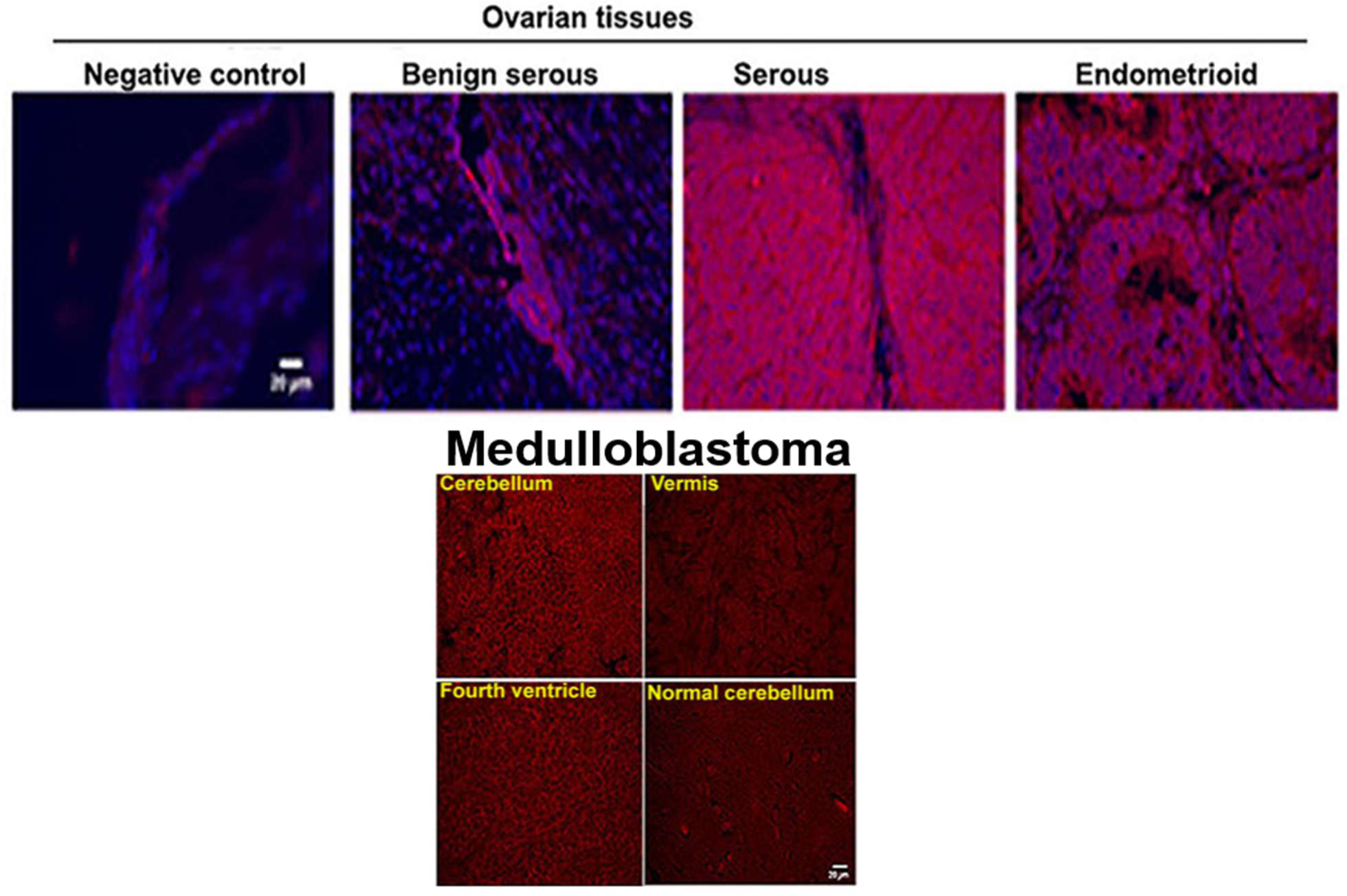
(**Upper**): Ovarian cancer tissues showed increased VDR expression compared to benign ovarian tissues. The cells were stained with primary VDR antibody and subsequently with corresponding secondary DyLight-594 (Vector Laboratories, Catalog number: DL2594). The micron bar (20μm) is shown. (**Lower**): Medulloblastoma tumor microarray (US Biomax Inc.) was stained with VDR primary and matched DyLight-594(Vector Laboratories, Catalog number: DI2594). Compared to normal cerebellum, malignant cerebellum and fourth ventricle tissues showed increased VDR expression. The micron bar (20μm) is shown.

**Supplementary Figure 4:**
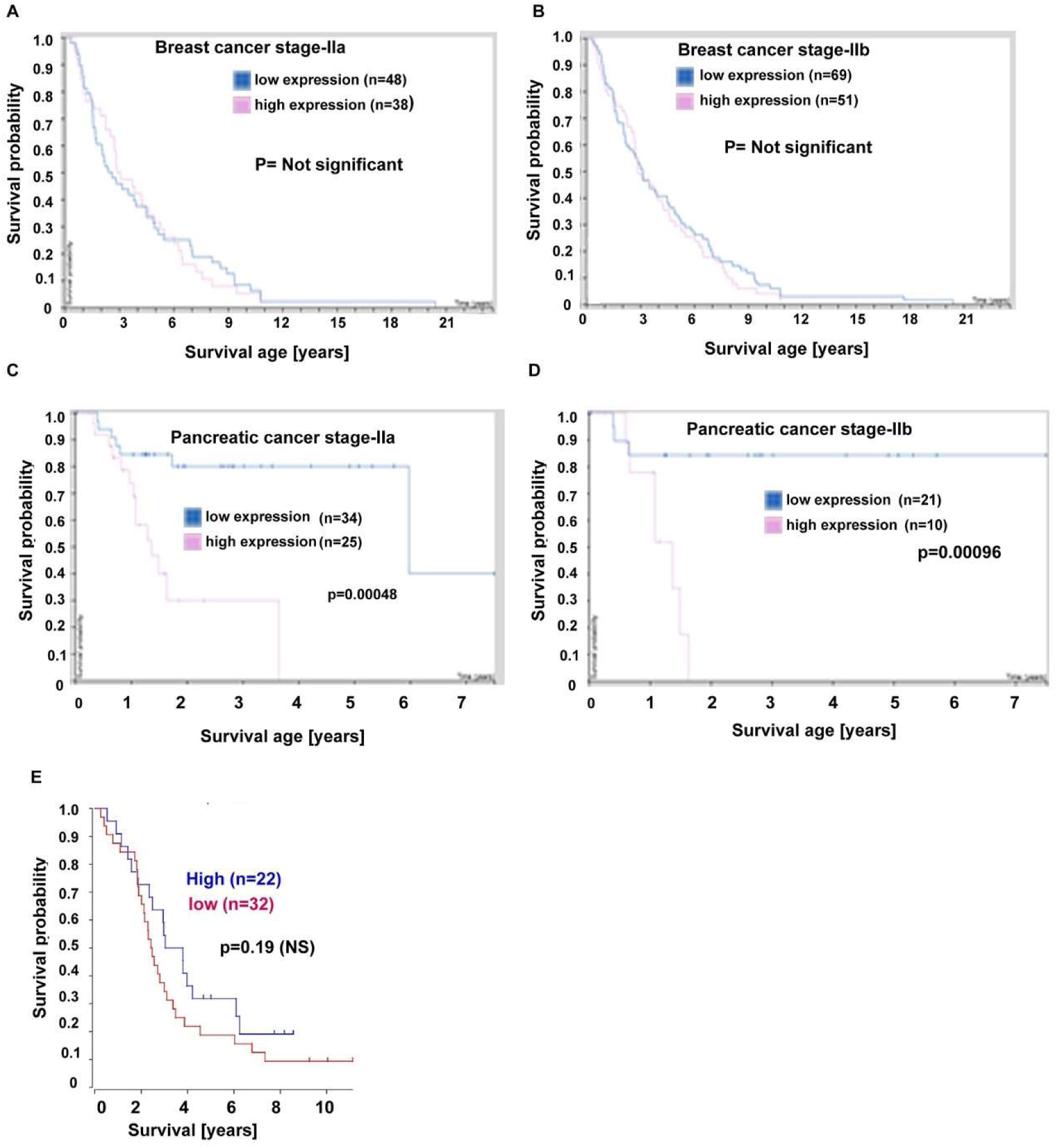
Impact of VDR mRNA enrichment on survival probability stratified by cancer stage. (**A-B**): Among stage-IIa and IIb breast cancer patients, VDR mRNA enrichment was not associated with increased mortalities. (**C-D**). Among stage-IIa and-IIb pancreatic patients VDR mRNA enrichment correlates significantly with increased mortalities. (**E**): Among ovarian cancer patients, disease stage (II) did not exhibit association of VDR mRNA enrichment with increased mortalities.

**Supplementary Figure 5:**
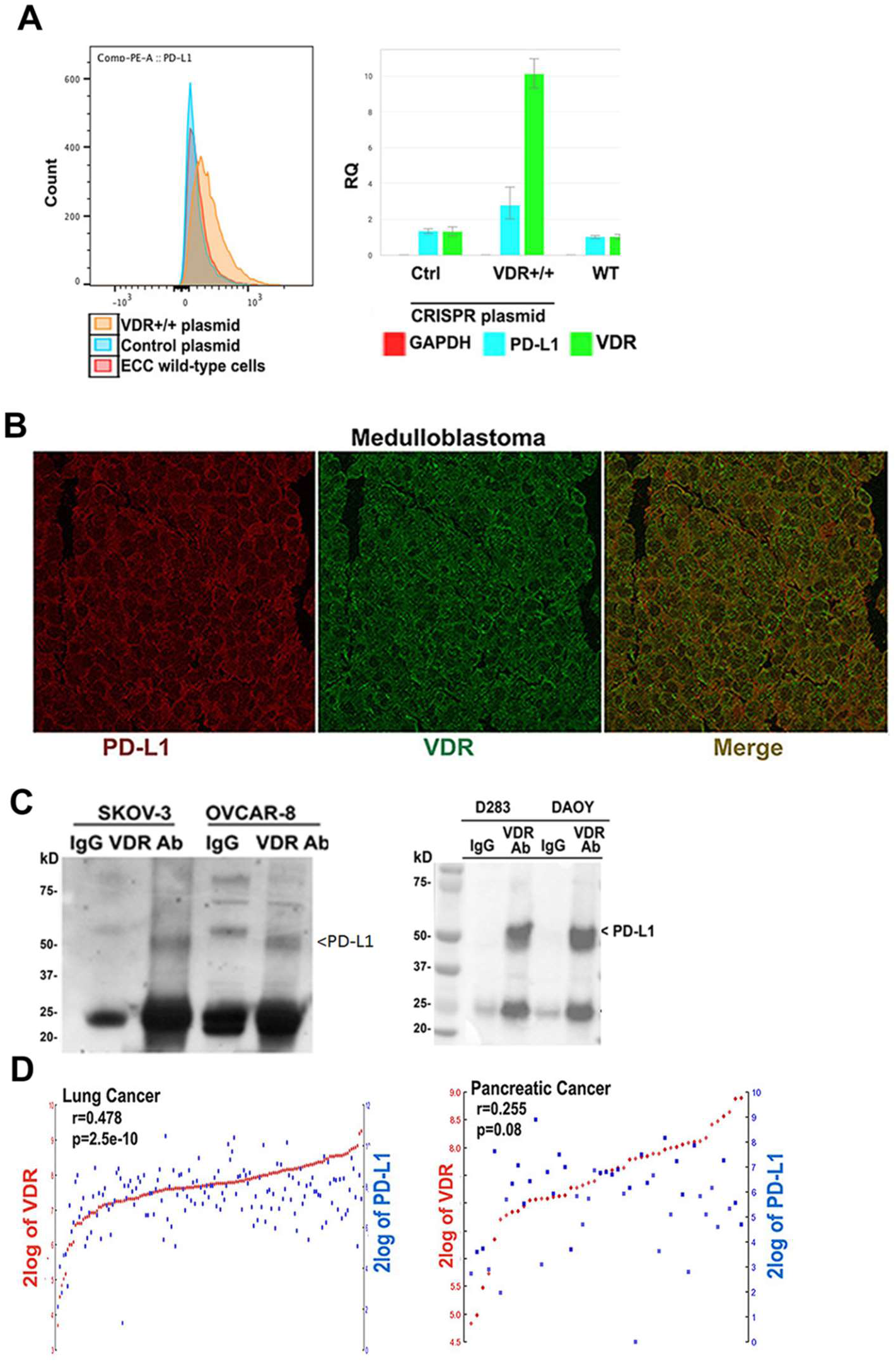
(**A**): Flow cytometric analysis showed that CRISPR plasmid mediated VDR upregulation resulted in increased surface expression of PD-L1 on ECC-1 endometrial cancer cells. VDR overexpressing plasmid and null vector transfected ECC-1 cells were analyzed by flow cytometry using PD-L1 antibody (PDL1-BV605, Biolegend, catalog no: 124321). Anti-mouse IgG Kappa beads (BD Biosciences, catalog number: 552843) were used as the negative control (**B**): VDR and PD-L1 exhibited co-localization in medulloblastoma tissues. Medulloblastoma tissues (US Biomax Inc, MD, USA) were processed, fixed and stained overnight with VDR primary antibody (Santa Cruz Biotechnology, catalog:SC-9164, dilution: 1:500). Slides were washed (PBST, 2×10mL, 5 minutes each) and stained with source matched secondary (DyLight-488, Vector Laboratories, catalog number: DL-1488) for an hour in dark. Tissues were washed (5x 10mL, 5 minutes each) and stained again with PD-L1 (Cell Signaling Technology, Mouse mAb catalog number: 29122, dilution: 1:1000) followed by staining with mouse secondary DyLight-594 antibody. Slides were bathed in PBST (5×10mL, 5 minutes each). DAPI containing mounting medium (Vectashield, Vector Laboratories, catalog number: H-1200) was applied and cover-slipped. Confocal images were recorded as described in the material and methods section. Micron bars =10μm. (**C**): A monoclonal VDR antibody (Santa Cruz Biotechnology, catalog number: H-81; SC-9164) immuno-precipitated PD-L1 from the total cell-lysates of SKOV-3 and OVCAR-3 cells. Similarly, the VDR antibody (Santa Cruz Biotechnology, catalog number: H-81; SC-9164) immuno-precipitated PD-L1 from the total cell lysates of DAOY and D283 medulloblastoma cells. (**D-H**): Two gene correlation analyses showed correlation between VDR and PD-L1 in lung (r=0.478, p=2.5e-10) and pancreatic cancer tissues (r=0.255, p=0.08). Publicly available lung and pancreatic cancer microarray data deposited at R2-Genomics Analysis and Visualization Platform were analyzed using their two-gene correlation analysis tool.

**Supplementary Figure 6:**
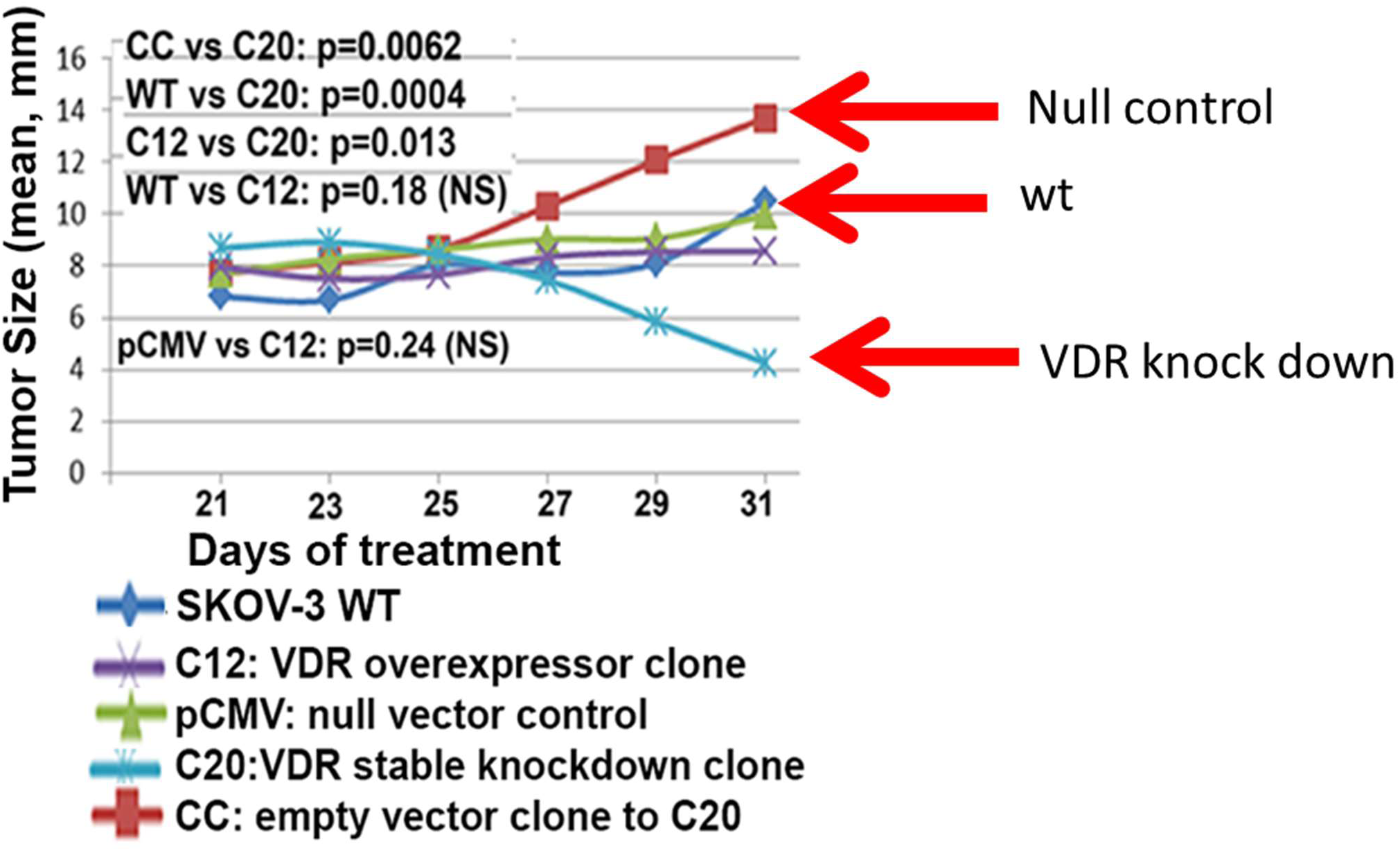
Stably VDR knock-down SKOV-3 cell clones (C20) showed negative tumor growth and burden during 31 days of tumor burden monitoring compared to wild-type SKOV-3, CC scrambled control, VDR high expressor clones (C12) and pCMV null vector transfected SKOV-3 cell clones. 1 million cells of wild-type and stably VDR overexpressing and under-expressing clones and their control counterparts were inoculated in nude mice. The tumor growth was monitored via measuring the longest diameter. Reduced tumor growth shown by clone-C20 (VDR knock-down) provided the key rationale that inhibiting VDR may reduce tumor burden.

### Supplementary Information-7a

**Method of Synthesis:** 7DHC (Sigma Aldrich) was stirred with N-methyl-1,2-4-triazolinedione (0.1:0.1 molar ratio) in dichloromethane at 0°C. Within four hours the pink color was found to be disappeared. The separated product (adduct) was filtered and coupled with bromoacetic acid in presence of DCC and catalytic amount of DMAP. Dichloromethane was removed using Buchi rotavapor and the crude product obtained was purified using a preparative thin layer chromatography plate. The band containing the product was collected and the compound was stripped off the silica gel by washing through MeOH:DCM (9:1). The solvent was removed using rotary evaporator and the compound (MeTC7) was collected after drying under vacuum as an off-white powder. The compound was stored at −20^0^C till used.

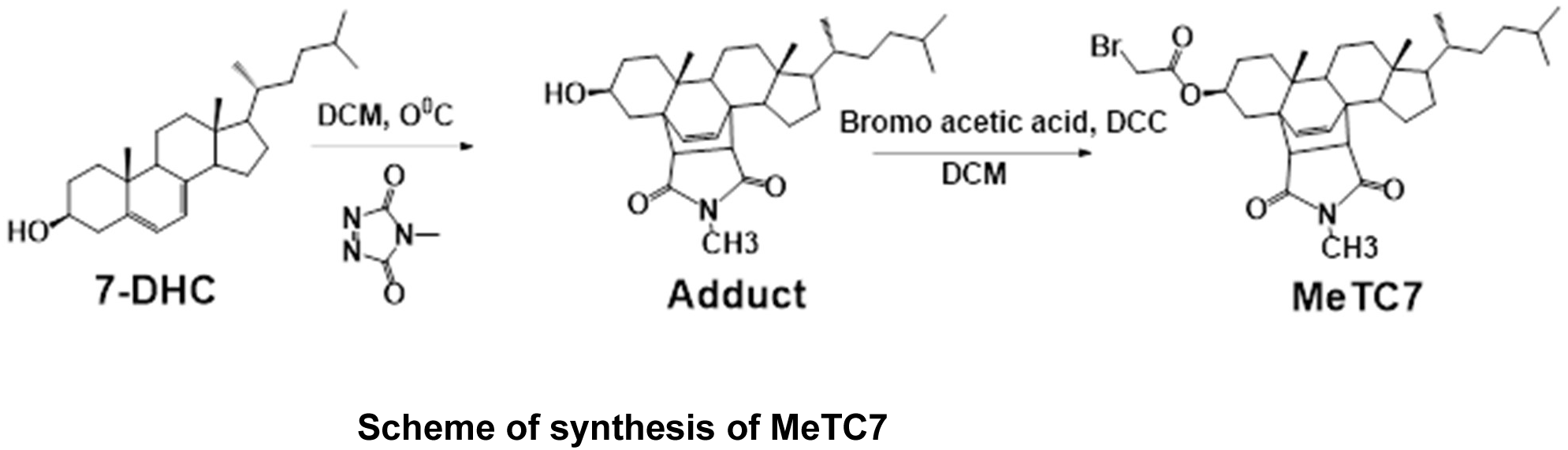

### Supplementary Data-7B

1. HNMR and 13C NMR data
2. H-1H COSY data

**Figure-7B-1:**
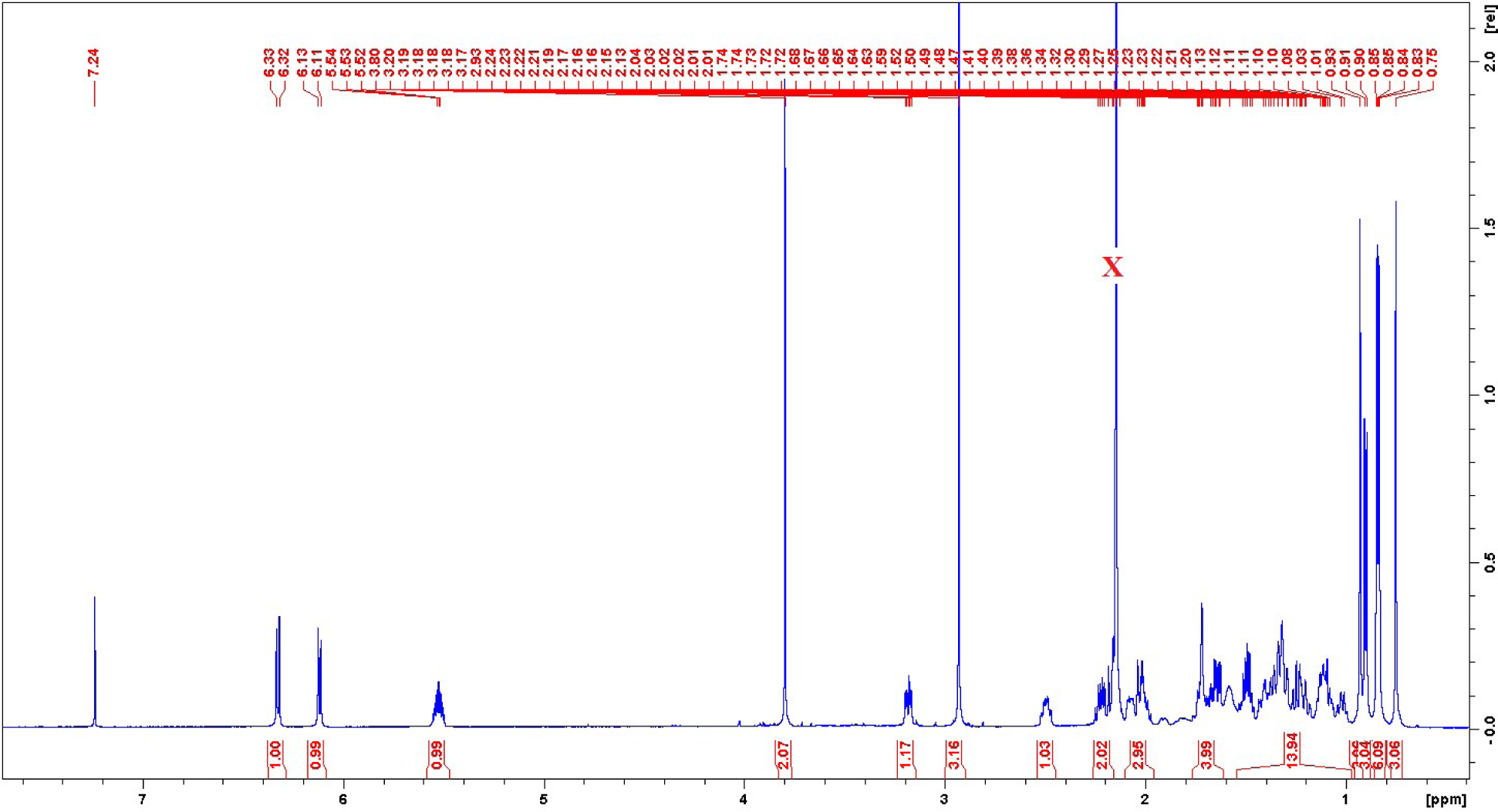
1H NMR profile of MeTC7

**Figure-7B-2:**
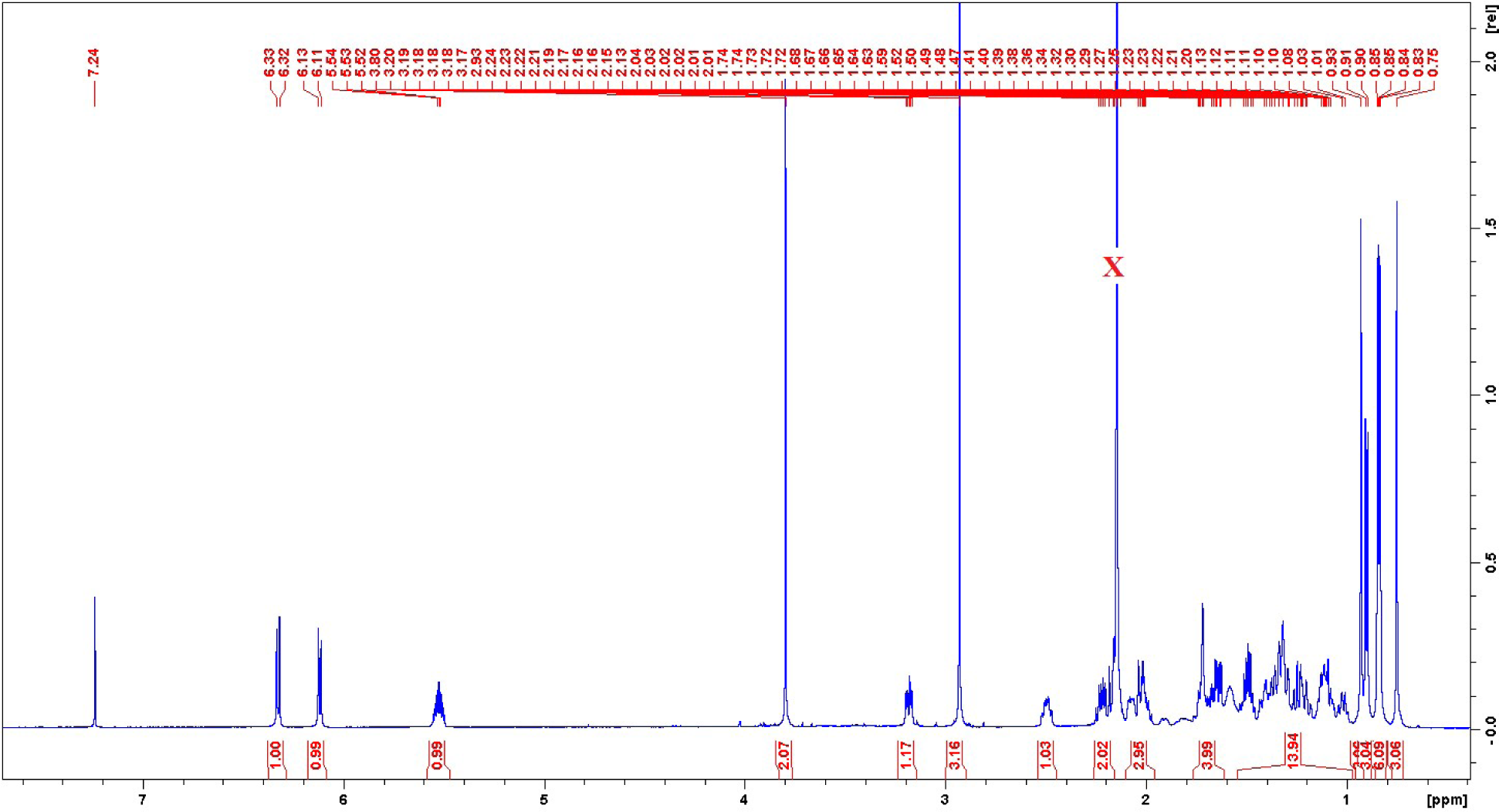
^13^C NMR profile of MeTC7

**Figure-7B-3:**
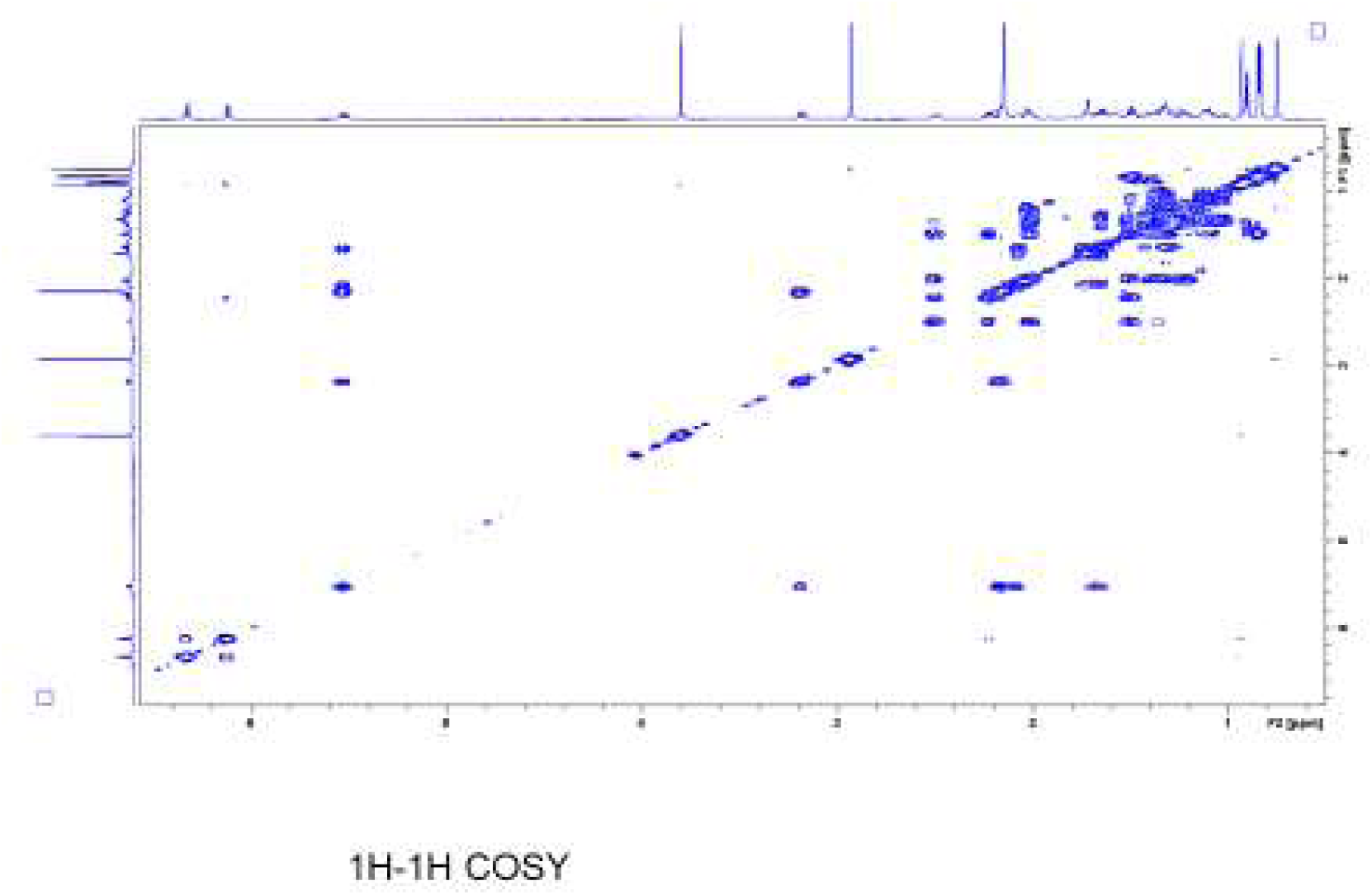
1H-1H NMR spectral profile of MeTC7

**Supplementary Information-8.**
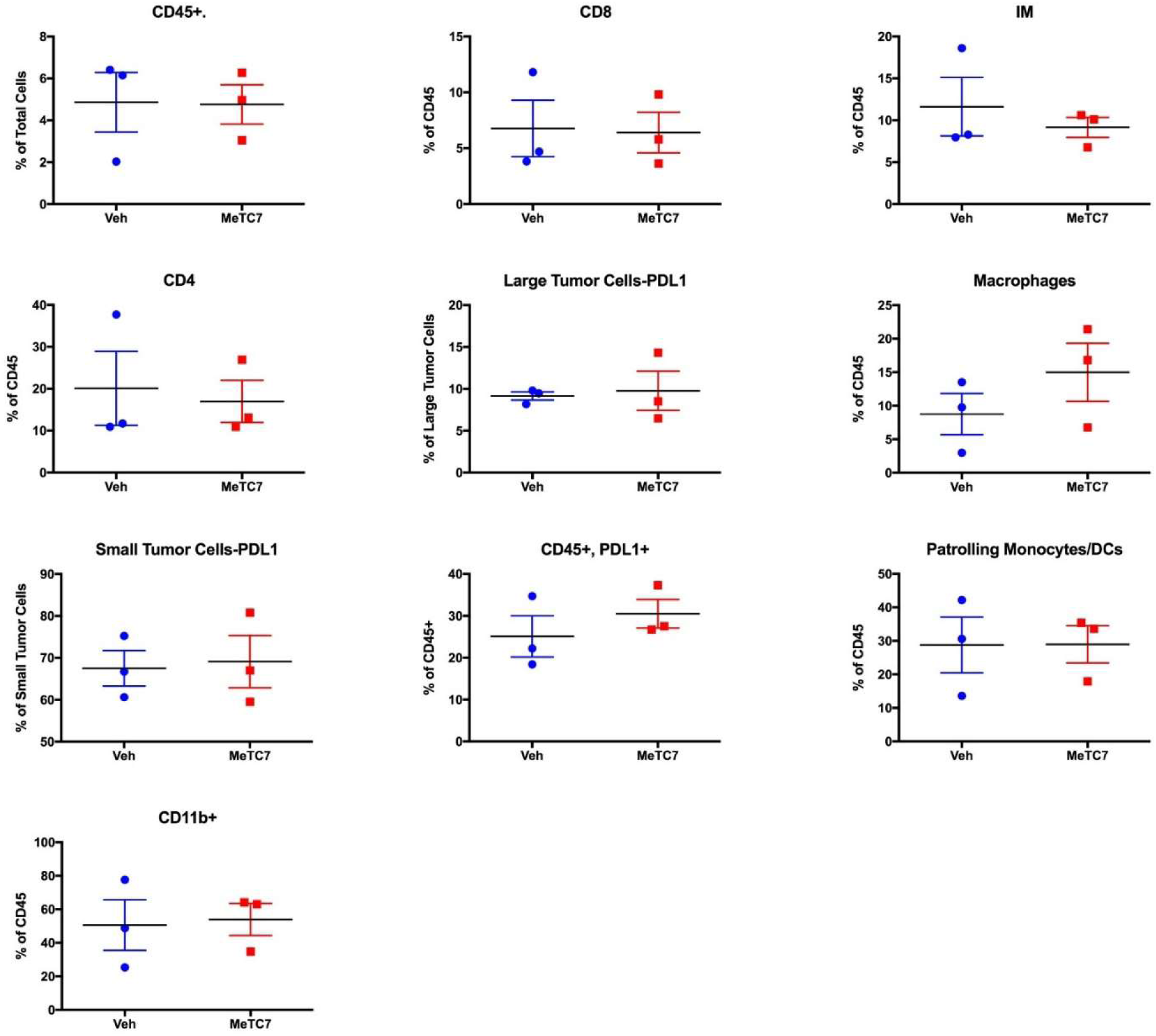
Spontaneous THMYCN tumor treated with vehicle or MeTC7 (10mg/kg, IP, once-daily) were harvested, broken into single cell suspension as described in materials and method section. The cells were stained with mouse flowcytometric antibodies representing CD45, PD-L1, CD11b, CD4 and CD8 antigens and those representing patrolling monocytes/DCs, macrophages and inflammatory monocytes. Analysis of the data showed that MeTC7 did not affect expression levels of immune markers analyzed.

## Supplementary Information-9

List and catalog numbers of the antibodies or primers used for the flow cytometric analyses and the ChIP assay.

1. CD45-FITC, BD Bioscience, cat#553080
2. CD8-PerCP-CyTM5.5, BD Bioscience, cat#551162
3. CD69-APC, BD Bioscience, cat#560689
4. PD1-PE-Cy7, Biolegend, cat#109110
5. PDL1-BV605, Biolegend, cat#124321
6. VDRE (f) primer sequence: 5’ GGA AAGGCAAACAACGAAGA-3’; VDRE(r): 5’GCGCTGAACTTCTAGGTGCT-3’

## Abbreviations

CDs: Cell differentiation markers (such as CD3, −8, −45 and −69)
irAEs: Immune related adverse reactions
MeTC7: Name of the drug candidate
MTAD: N-methyl-1,2,4-triazoline dione
PTAD: N-phenyl-1,2,4-triazoline dione
RT: Radiation Therapy
PD-L1: Programmed death receptor ligand
PD-1: Programmed death receptor
VDR: Vitamin-D receptor

